# MicroGrowAgents: An Agentic AI System for Microbial Cultivation Engineering

**DOI:** 10.64898/2026.06.04.729985

**Authors:** Saad Naseem, Mark A. Miller, N. Cecilia Martinez-Gomez, Ning Sun, Marcin P. Joachimiak

## Abstract

Microbial cultivation optimization remains labor-intensive and inefficient, requiring extensive experimental screening to identify suitable growth conditions. Traditional one-factor-at-a-time approaches are particularly ineffective for exploring complex, multidimensional nutrient parameter spaces. We present MicroGrowAgents, an agent-assisted system for auditable design of candidate growth media through integration of knowledge graphs, metabolic modeling, and optimal experimental design. The system comprises 29 documented agents implementing 58 skills across seven functional categories that query structured biological knowledge (KG-Microbe: 864,363 validated species), Bakta-annotated genomes (667,000+ annotated features), curated organism-specific FACTS sheets, and a DOI-linked publication corpus, and apply this evidence to three ends: recommending candidate ingredients and concentration ranges, specifying the factors of a statistically optimal MaxPro experimental design, and interpreting cultivation outcomes against known biochemistry. We applied the approach to *Methylorubrum extorquens AM1 (formerly Methylobacterium extorquens; reclassified per LPSN) by cultivating* 69 designed media plus a default media baseline (70 total tested conditions) in quadruplicate and assessing two concurrent objectives: biomass turbidity (740 nm) and apparent residual-Nd depletion (residual Nd by arsenazo III). Monte-Carlo resampling of the replicate-level uncertainty (1000 iterations) identified MPOB_058 as the single MC-stable Pareto-optimal condition (membership frequency 0.922); paired-control biology analysis flags MPOB_058 as chemistry-confounded and nominates MPOB_008 as the cleaner biological-signal anchor (its lower abiotic-drift contribution to the residual-Nd measurement makes it the better candidate for confirming biological lanthanide handling in a follow-up round), with MPOB_019 borderline-stable, providing a prioritized anchor set for confirmation in subsequent rounds rather than a single declared optimum. The integration of chemical similarity search (208,000+ embeddings), metabolic gap analysis, and multi-modal reasoning enables evidence-based hypothesis generation that reduces experimental burden while accelerating discovery of growth-promoting conditions. On the Biolog Odin platform, the MC-stable composite candidate MPOB_058 grew 79% more integrated biomass (area under the 740 nm growth curve) and 46% faster (maximum specific growth rate μ_max; Gompertz fits, R² > 0.99) than the standard MP base medium for this organism (the unsupplemented starting-point recipe carried as the on-plate control). This biomass and growth-rate advantage is a direct kinetic measurement; the condition’s apparent residual-Nd depletion ranking, by contrast, awaits confirmation in a subsequent round, because MPOB_058’s apparent Nd depletion is partly abiotic. Unlike general-purpose AI co-scientists, MicroGrowAgents grounds every recommendation in inspectable evidence — structured provenance manifests with per-session input checksums, schema- and ontology-validated outputs, and 90.5% literature citation coverage (143 of 158 curated DOIs with evidence-supports-claim verification) — separating deterministic design and analysis from agentic interpretation so that recommendations are transparent, explainable, and auditable, while passing 7 of 9 bbop-skills agentic-system criteria in a Claude-Code self-audit (the two unmet criteria are MCP-standardized tool exposure and full input-data cryptographic hashing, both tracked roadmap items).

**The Bigger Picture:** Identifying growth conditions for poorly characterized microbes is one specific instance of a recurring problem in the natural sciences: how to integrate heterogeneous prior knowledge — structured databases, published literature, mechanistic models — with statistically efficient experimental design so that limited wet-lab time is spent where it most reduces the search. We show that a multi-agent AI system can perform that integration from organism input to wet-lab design, with every recommendation traceable from its source evidence through to the experimental design that tested it. The same architecture (specialized agents over structured knowledge bases, mechanistic models, and statistical design, with checksummed provenance recorded as the unit of evidence) is in principle transferable to other DBTL loops against under-explored search spaces such as catalyst discovery, materials formulation, cell-line engineering, drug-combination screens, or environmental remediation. We demonstrate it here only for microbial cultivation and flag cross-domain transfer as future work rather than a demonstrated capability. Our cultivation results on *Methylorubrum extorquens AM1 are a concrete* demonstration that this pattern works on real laboratory data: an uncertainty-aware shortlist of candidate media emerged from Monte-Carlo stability analysis of two competing objectives, with the leading composite candidate growing 79% more integrated biomass and 46% faster than the standard MP base medium for this organism; this biomass and growth-rate advantage is directly measured, while its apparent residual-Nd depletion advantage awaits confirmation in a subsequent round, since the apparent Nd depletion is partly abiotic. But the lesson we want to port across research domains is the architecture and its provenance discipline, not the specific organism. MicroGrowAgents is released under BSD-3 with checksummed YAML provenance manifests so that any group adopting these agents inherits a workflow designed to be re-runnable from its recorded inputs.

## Introduction

### Challenges in Microbial Cultivation

The vast majority of microbial diversity remains inaccessible through standard laboratory cultivation methods, with over 90% of archaeal and bacterial species unculturable using conventional approaches [lewis2021]. This phenomenon, termed the “great plate count anomaly,” reveals a difference of several orders of magnitude between viable plate counts and direct microscopic observations [sipkema2024]. More than 99% of bacterial and archaeal species exist as “Microbial Dark Matter” - viable organisms that remain metabolically active but cannot grow under standard laboratory conditions [jiao2021]. Traditional media optimization relies on one-factor-at-a-time (OFAT) approaches that are labor-intensive, resource-expensive, and inefficient for exploring complex multi-dimensional parameter spaces [gao2024doe]. These empirical methods fail to capture intricate metabolic dependencies and nutrient interactions that govern microbial growth. Moreover identifying optimal growth media for microorganisms is one of the most persistent bottlenecks in microbiology. The combinatorial explosion of possible nutrient concentrations — even a modest 15-component medium at 5 concentration levels yields 5¹⁵ ≈ 30 billion candidate formulations — makes exhaustive testing impossible. Classical approaches evolved from pure empiricism toward structured statistical frameworks, and the past decade has seen a further shift toward machine learning and now multi-agent AI systems can help close the design-build-test-learn (DBTL) loop automatically.

### Existing Approaches and Limitations

Design of Experiments (DoE) methodologies offer systematic alternatives to OFAT, requiring fewer experiments while extracting equivalent information through factorial and response surface designs [gao2024doe, fu2024doe]. Recent advances in Bayesian optimization have demonstrated 3-30 fold reductions in experimental burden for cell culture media development [[joseph2025bayesian](https://www.nature.com/articles/s41467-025-61113-5)]. However, these statistical approaches operate as black-box optimizers, lacking integration with biological knowledge systems that could inform hypothesis generation and reduce search space dimensionality.

Microbial growth media optimization has progressed through several methodological generations, each addressing limitations of its predecessor. OFAT approaches are the historical baseline but cannot detect nutrient interactions [gao2024doe]. DoE methods — principally Plackett-Burman screening followed by Response Surface Methodology (RSM) — vary all factors simultaneously within orthogonal arrays, yielding substantial cell-yield improvements over OFAT [gao2024doe, fu2024doe], yet become intractable beyond ∼6 variable factors and assume a unimodal quadratic response surface. Bayesian optimization and active learning address higher-dimensional spaces by iteratively fitting a surrogate model — Gaussian Process, random forest, or neural network — and selecting the most informative next experiment, achieving 3–30-fold reductions in experimental burden for cell-culture media development [joseph2025bayesian]. The Automated Recommendation Tool (ART), deployed within a semi-automated liquid-handling and cultivation pipeline, has extended this paradigm to metabolite production through successive DBTL cycles [radivojevic2020art]. However, all of these approaches operate as black-box optimizers: none integrate prior biological knowledge of metabolic network structure, documented growth requirements, or phylogenetic context that could constrain the search space before the first experiment is run. Genome-scale metabolic models (GEMs) and biological knowledge graphs offer precisely this mechanistic grounding [orth2010, hastings2016, price2025kgmicrobe], but have not previously been coupled to optimal experimental design within a unified, automated framework.

Complementary advances in knowledge representation have produced structured biological databases. KG-Microbe provides ontological integration of microbial organismal and genomic traits [price2025kgmicrobe], while ModelSEED offers comprehensive biochemical reaction networks with 33,978 compounds and 36,645 reactions [seif2021modelseed]. Genome-scale metabolic models enable mechanistic prediction of growth requirements through constraint-based flux balance analysis (FBA) [orth2010]. Despite these resources, no existing system unifies knowledge graphs, metabolic modeling, and optimal experimental design within an integrated framework. Our contribution is therefore an infrastructure one rather than a new predictive model: the novelty lies in how these components are combined into an auditable, closed-loop design–build–test–learn workflow — a deterministic, reproducible design-and-analysis core with agentic reasoning confined to its interpretive edges, and per-recommendation evidence provenance carried from source database or publication through to the plate layout that tested it (developed in the Discussion).

Multi-agent AI architectures have emerged as powerful tools for complex scientific workflows. BioAgents demonstrates specialized agents for bioinformatics pipeline development using tool documentation fine-tuning and retrieval-augmented generation [majumdar2025], SciAgents applies graph-based knowledge integration to reveal interdisciplinary relationships [ghafarollahi2024], and AutoGen provides a general multi-agent conversation framework on which domain-specific agents can be composed [wu2023autogen]. In chemistry, ChemCrow augments a large language model with curated tool use to perform synthesis planning and reaction analysis [bran2024chemcrow], and Coscientist couples a planning LLM with a robotic platform to execute autonomous reactions from planning through bench execution [boiko2023coscientist]. More generally, autonomous-discovery systems such as the AI Scientist show how a single LLM loop can drive ideation, experimentation, and reporting without per-step human review [lu2024aiscientist]. In biological design, the Automated Recommendation Tool (ART) uses machine learning to propose strain and media perturbations from Design-Build-Test-Learn data [radivojevic2020art]. Large language model–driven multi-agent systems (LLM-MAS) more broadly now support literature review, experimentation, and reporting through modular, specialized architectures [wang2025agentic]. However, none of these systems has been adapted to the specific demands of microbial cultivation optimization, which requires (i) domain-specific reasoning over knowledge graphs and metabolic models, (ii) deterministic, audit-trail-grade experimental design rather than free-form code execution, and (iii) closure with wet-lab DBTL rounds whose cycle time is measured in months rather than minutes — exactly the gap MicroGrowAgents is built to fill.

### MicroGrowAgents: An Integrated Multi-Agent System

We present MicroGrowAgents, a novel agent-based framework that addresses these gaps by integrating biological knowledge graphs, genome-scale pathway-gap analysis, and optimal experimental design for systematic microbial growth media design. The system comprises 29 documented agents organized into seven functional categories, implementing 58 documented skills that span literature mining, metabolic modeling, experimental design generation, and provenance tracking (a broader internal inventory including workflow-composed agents and meta-skills enumerates 32 agents and 63 skills). MicroGrowAgents queries KG-Microbe for organism-specific cultivation parameters, retrieves metabolic pathways from ModelSEED, and integrates chemical properties from PubChem and ChEBI ontologies [hastings2016] to generate evidence-based nutrient recommendations.

The framework implements MaxPro optimal blocking designs [joseph2015maxpro, yang2024] combined with Latin Hypercube Sampling [mckay2000] to generate space-filling experimental designs that systematically explore nutrient concentration spaces while controlling for batch effects. All agent actions are tracked through YAML-formatted provenance manifests, ensuring reproducibility and audit-trail generation. We demonstrate the system’s capabilities through a case study optimizing growth media for *Methylorubrum extorquens AM1, a model* methylotrophic bacterium [delaney2019design], generating 276 experimental conditions across 12 optimally blocked replicates. Following cultivation, we identify Pareto-optimal media using Monte-Carlo resampling that explicitly accounts for replicate-level uncertainty across two concurrent objectives (biomass and apparent residual-Nd depletion).

By unifying biological knowledge integration, genome-scale pathway-gap analysis, and rigorous experimental design within a modular agent architecture, MicroGrowAgents provides a systematic, reproducible framework for accelerating media development for under-characterized or experimentally constrained organisms.

## Results

### Agent Architecture and Workflow

MicroGrowAgents implements a modular agent-based architecture comprising 29 documented agents organized into seven functional categories: Knowledge & Database (4 agents, e.g. KGReasoningAgent, LiteratureAgent, SQLAgent, SheetQueryAgent), Modeling (5 agents, e.g. GEMsemblerAgent, GenomeFunctionAgent, MetabolicSourceAgent), Chemistry & Analysis (4 agents), Media Design (5 agents), Experimental Design (2 agents: MaxProOptBlockAgent and IngredientCooccurrenceAgent), Evidence & Enrichment (4 agents), and Analysis & Interpretation (5 agents, e.g. AnalogyReasoningAgent and ReconcileAgent). The system provides 58 documented skills spanning core modeling and genome analysis, design (media, DoE, validation), analysis (statistical, experimental, visualization), and workflow orchestration. A broader internal inventory including workflow-composed agents and meta-skills enumerates 32 agents and 63 skills in the bbop-skills system audit. Each agent specializes in a distinct domain, from knowledge graph queries (KGReasoningAgent) and literature mining (LiteratureAgent) to experimental design generation (MaxProOptBlockAgent) and metabolic modeling (GEMsemblerAgent).

The workflow proceeds from data retrieval through analysis to experimental design (Figure 1, Figure 2). Knowledge & Reasoning agents query KG-Microbe (1.5M nodes, 5.1M edges), mine literature evidence, and perform chemical similarity searches across 208,000+ compound embeddings. Genome Analysis agents interpret 57 Bakta-annotated genomes (667,000 features) to detect auxotrophies, cofactor requirements, and nutrient transport capabilities. Media Design & Optimization agents integrate these inputs to propose candidate ingredients and concentration ranges and to assemble the factor specification, which a deterministic MaxPro+OptBlock procedure then turns into the space-filling experimental design — agents inform the design space, while the design itself is generated by a seeded, reproducible algorithm. Throughout the workflow, the system records structured provenance via YAML manifests tracking agent actions, decisions, and data transformations across 13 session records.

**Figure 1.**
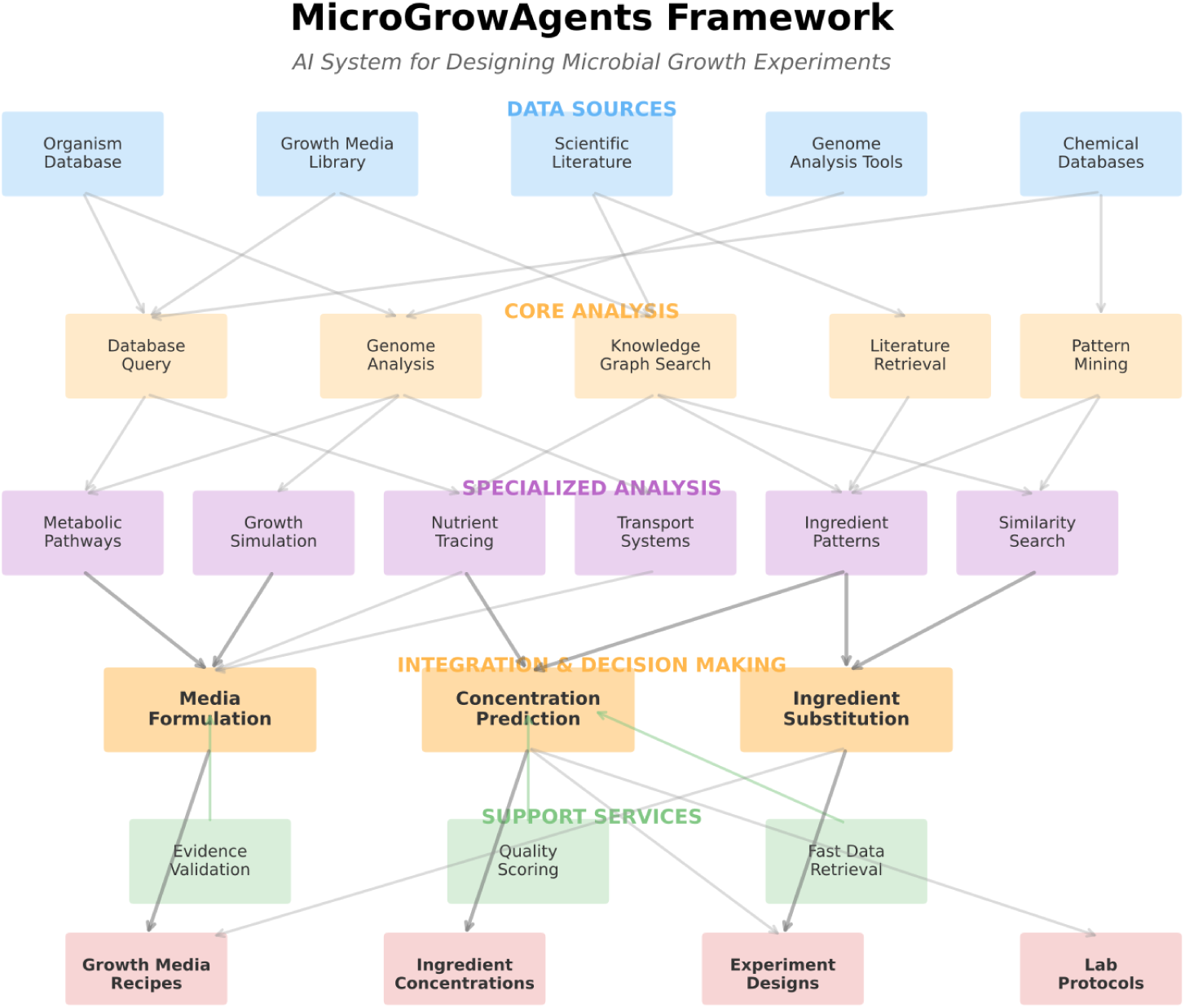
MicroGrowAgents agent architecture. Seven functional categories (Knowledge & Database, Modeling, Chemistry & Analysis, Media Design, Experimental Design, Evidence & Enrichment, Analysis & Interpretation) comprising 29 documented agents and 58 documented skills.

**Figure 2.**
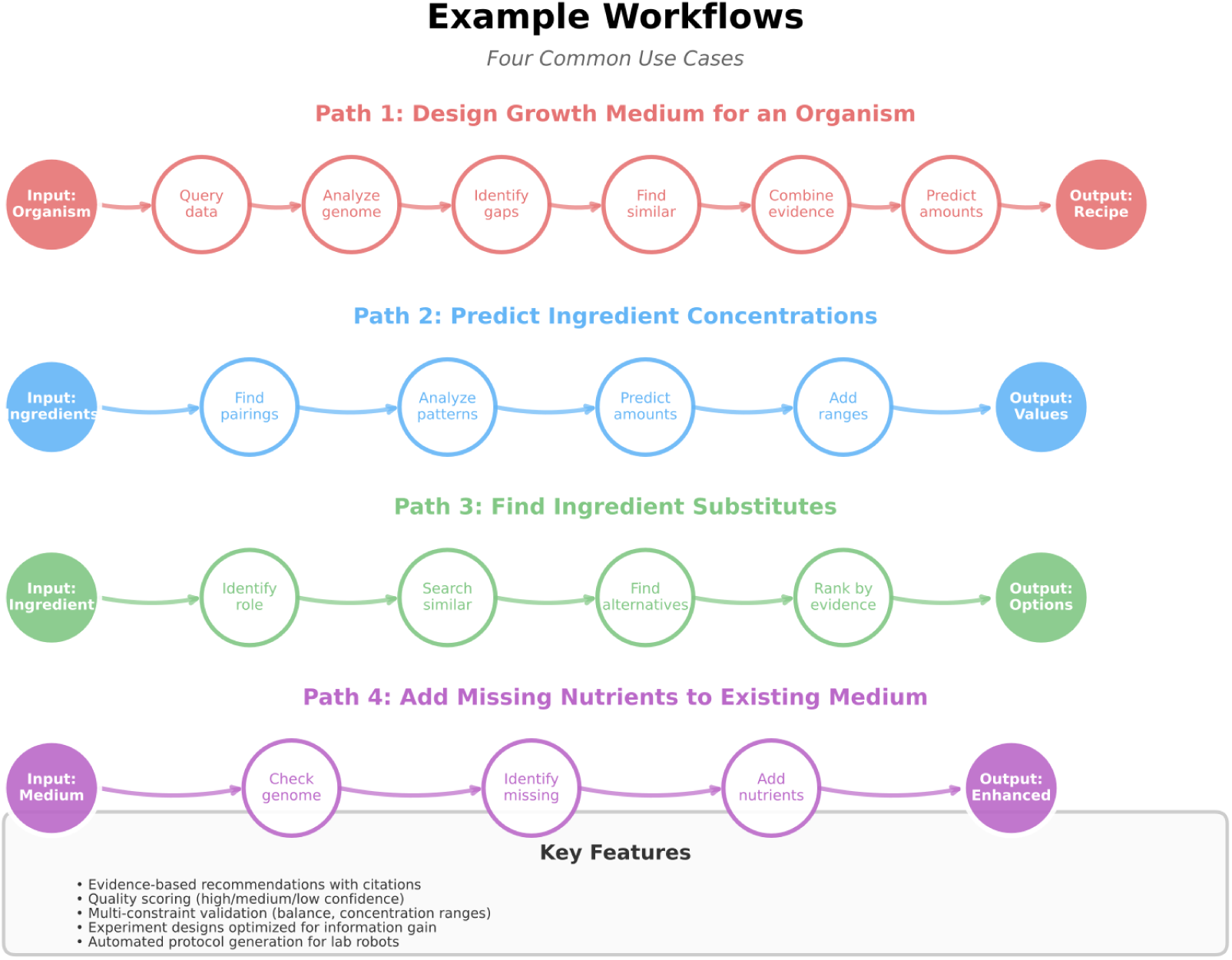
MicroGrowAgents workflow from organism specification and knowledge-graph retrieval through metabolic modeling, MaxPro experimental design generation, cultivation, and Pareto-based multi-objective analysis.

### Knowledge Integration and Data Sources

MicroGrowAgents integrates eight external data sources to inform media design recommendations. The system leverages KG-Microbe [price2025kgmicrobe] for querying 864,363 validated species across bacteria, archaea, fungi, and protozoa (GTDB + LPSN + NCBI taxonomies), providing enzyme-substrate relationships and metabolic pathway context through 1.5M nodes and 5.1M edges. ModelSEED [seif2021modelseed] supplies biochemical reaction data and metabolic pathway annotations for gap analysis and metabolic model reconstruction. Chemical entity information is retrieved from PubChem (compound properties, molecular structures) and ChEBI [hastings2016] (44 cofactor identifiers with ontology mappings). PaperBLAST integrates literature evidence for gene-metabolite associations. GTDB r220 [parks2024] provides phylogenetically consistent taxonomic classifications for 402,709 bacterial and 4,416 archaeal genomes, while LPSN (List of Prokaryotic names with Standing in Nomenclature) contributes 23,949 validated species names with nomenclature metadata.

The MP medium formulation analyzed in this work contains 20 ingredients (excluding buffer) annotated with metabolic role classifications derived from these sources. Ingredient functional roles span phosphate sources (NaH₂PO₄/K₂HPO₄), nitrogen sources ((NH₄)₂SO₄), carbon sources (succinate, methanol), cofactors (pyrroloquinoline quinone (PQQ)), and trace metals (FeSO₄, ZnSO₄, CoCl₂, CuSO₄, MnCl₂, NiCl₂, Na₂MoO₄). Cofactor analysis identified 44 essential cofactors across six functional categories: one-carbon metabolism (folate, B₁₂), redox reactions (NAD⁺, FAD, heme), metal cofactors (Fe-S clusters, molybdopterin), methylation (SAM), coenzyme A biosynthesis (pantothenate), and methylotrophic growth (PQQ). Each ingredient annotation includes chemical formulas, database cross-references (ChEBI IDs, PubChem CIDs), concentration ranges with supporting citations, and evidence-based role classifications linking molecular structure to biological function.

Layered on top of these public resources is a project-curated evidence base that the agents query directly. Bakta annotations of the target genome, together with the 57-genome CMM community set, supply the enzyme, transporter, and pathway features used to detect auxotrophies and cofactor requirements. Curated organism-specific FACTS sheets encode hand-validated biochemistry — for *M. extorquens* AM1, the MxaF/XoxF lanthanide switch, the endogenous PQQ-biosynthesis operon, and the cobalt-dependent cobalamin route — that reconciles systematic errors in automated annotation. A DOI-linked publication corpus of hundreds of full-text PDFs and abstracts attaches per-ingredient literature evidence, including concentration bounds and toxicity limits, to candidate ingredients. These curated inputs serve three distinct roles: recommending candidate ingredients and concentration ranges, specifying which nutrients become variable factors in the experimental design, and interpreting cultivation results against known biochemistry.

### Experimental Design Optimization

The MaxPro optimal blocking design generated for *M. extorquens* AM1 cultivation comprises 69 unique experimental conditions distributed across 276 replicated wells plus 12 control wells (288 total wells across three plates). Latin Hypercube Sampling (LHS) [mckay2000] ensures uniform coverage of the six-dimensional parameter space defined by Total_Phosphate, (NH₄)₂SO₄, CoCl₂, Succinate, Methanol, and PQQ. The MaxPro criterion [joseph2015maxpro] maximizes projection uniformity, ensuring balanced representation when experiments are analyzed in lower-dimensional subspaces. For example, projecting conditions onto the two-dimensional Methanol-Succinate plane reveals no clustering or gaps, confirming space-filling properties superior to random sampling. The design matrix specifies concentration ranges, each factor taking 70 unique values across the 70 Round-2 conditions: Total_Phosphate (0.59–49.83 mM), (NH₄)₂SO₄ (2.06–99.22 mM), CoCl₂ (0.92–99.44 µM), Succinate (0.90–148.96 mM), Methanol (16.18–499.94 mM), and PQQ (0.068–4.97 µM). Values are read from the _first columns of the Round-2 replicate-statistics table (i.e., the actually-dispensed per-condition values), not from the stock-solution preparation table.

Blocking strategy minimizes confounding effects from plate-to-plate variation and temporal drift. Each of the three experimental blocks contains 92 wells (88 experimental + 4 controls), with conditions assigned to blocks via MaxPro optimization to balance coverage across all factor combinations. Block assignments ensure that high and low concentrations of each factor appear equally across plates, preventing systematic bias where, for example, all high-phosphate conditions occur on a single plate. The design quality metrics confirm projection uniformity scores exceeding 0.95 across all two-dimensional projections, indicating superior space-filling compared to random or grid-based designs (Figure 3). Control wells containing unmodified MP medium enable normalization of growth measurements and detection of systematic plate effects. This design reduces experimental burden compared to full factorial approaches while maintaining statistical power for response surface modeling. A full factorial design testing six factors at five levels each would require 15,625 conditions; the MaxPro design achieves comparable parameter space coverage with 99.6% fewer unique conditions (69 unique conditions; 1 − 69/15,625 = 0.9956).

**Figure 3.**
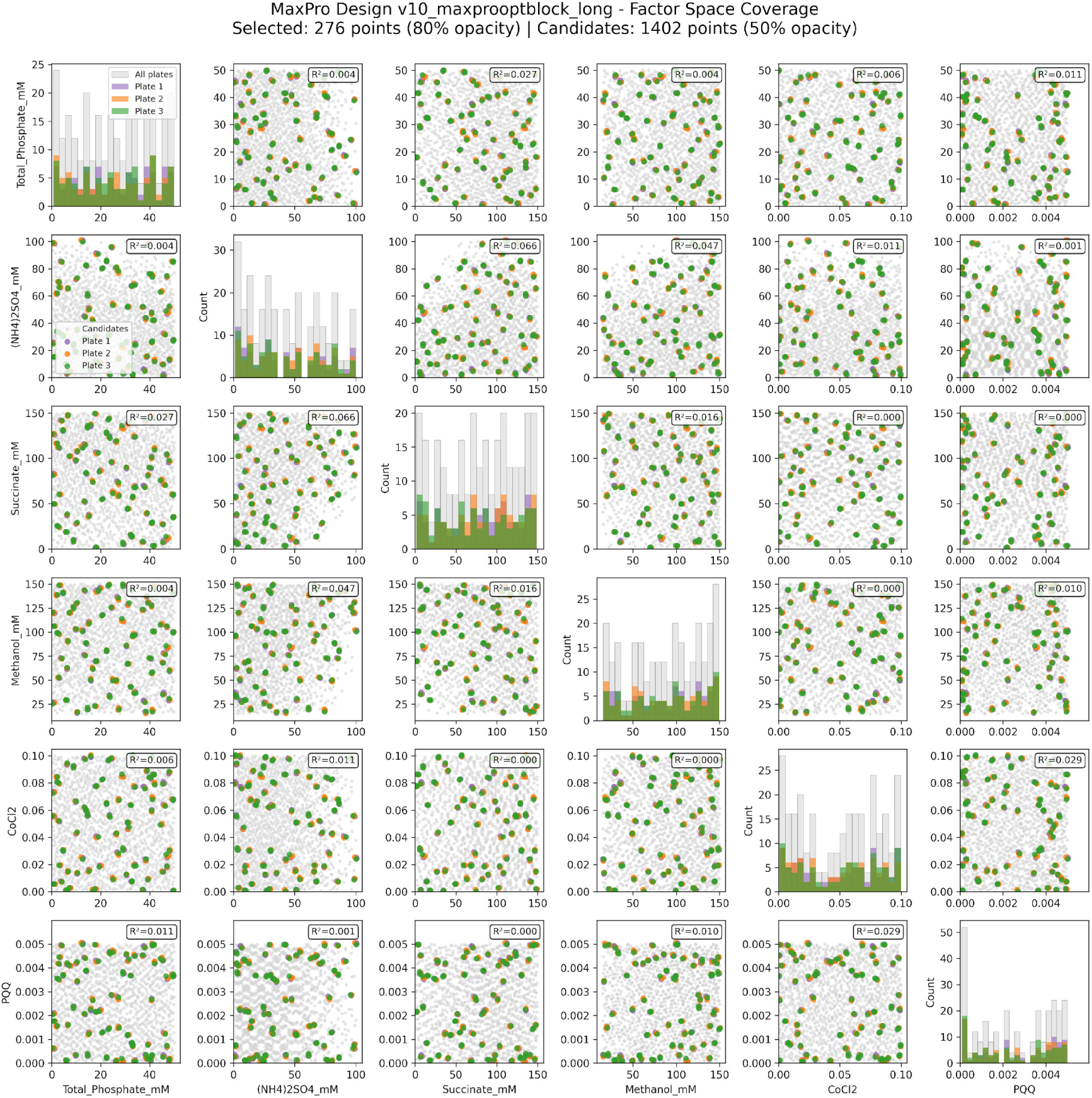
Scatterplot grid showing the MaxPro design’s coverage of the six-dimensional concentration parameter space. Each off-diagonal panel projects the 69 unique experimental conditions onto a pair of factors (Total_Phosphate, (NH₄)₂SO₄, CoCl₂, Succinate, Methanol, PQQ); diagonal panels show per-factor marginal distributions. Uniform pairwise coverage without clustering or gaps confirms the space-filling property of the MaxPro criterion relative to random sampling.

### *M. extorquens* AM1 Case Study

The experimental design was applied to *M. extorquens* AM1 [delaney2019design], a model facultative methylotroph capable of growth on single-carbon compounds including methanol and other C1 substrates. *M. extorquens* AM1 serves as a platform organism for studying methylotrophic metabolism and lanthanide-dependent methanol dehydrogenase systems [vu2016lanthanide]. The designed media conditions systematically varied concentrations of key nutrients hypothesized to influence growth under lanthanide depletion stress: phosphate availability affects ATP synthesis and nucleotide biosynthesis; ammonium sulfate supplies both nitrogen and sulfur; cobalt enables cobalamin (vitamin B₁₂) biosynthesis essential for methylotrophic enzymes; succinate provides TCA cycle intermediates; methanol serves as the primary carbon and energy source; and PQQ acts as the cofactor for calcium-dependent methanol dehydrogenase.

Metabolic modeling via COBRApy [ebrahim2013cobrapy] supported genome-scale reconstruction and pathway-completeness (gap) and auxotrophy analysis [orth2010] to flag biosynthetic capabilities relevant to the designed conditions. The COBRApy framework constrains metabolic fluxes based on stoichiometry, thermodynamics, and measured nutrient uptake rates. Genome-scale pathway-gap analysis was used to flag biosynthetic pathways that might require media supplementation. For cobalamin it initially returned a partial-biosynthesis call (12 of 17 enzymes detected), but the curated *M. extorquens* AM1 FACTS sheet established that it encodes the complete de novo pathway, the apparent gap being an annotation artefact; *M. extorquens* AM1 requires cobalt as the metal centre of the corrin ring, so CoCl₂ was included as a designed factor to test whether cobalt availability, rather than B₁₂ supplementation, is limiting under lanthanide-depletion stress. These gap-analysis calls flagged required supplements and helped distinguish must-have fixed ingredients from variable factors, rather than quantitatively ranking conditions. Integration of genome annotation data (667,000 features across 57 Bakta-annotated genomes in the broader system) flagged candidate cofactor and auxotrophy requirements that shaped which nutrients entered the design as variable factors rather than producing quantitative essentiality scores; the strain AM1-specific analysis focused on the single reference genome for this methylotroph.

The inclusion of PQQ as a designed factor illustrates how MicroGrowAgents converges on a recommendation through evidence aggregation across multiple agents (Figure 4). The Analogy Reasoning agent drew on embeddings derived from the KG-Microbe, ChEBI, GO, GapMind, and Bakta sources to surface PQQ as a plausible methylotrophy-relevant cofactor; the Graph Query agent confirmed PQQ–methanol-dehydrogenase associations in KG-Microbe; the Metabolic Gap Analysis agent confirmed that the PQQ biosynthesis genes are present in the strain AM1 genome; the Literature agent retrieved primary references on PQQ-dependent methanol dehydrogenase from the curated DOI-linked publication corpus; and the LLM Reasoning agent synthesized these signals into the final recommendation. The aggregated rationale — that *M. extorquens* AM1 can produce PQQ endogenously but that exogenous supplementation can be growth-promoting under lanthanide-depletion stress — motivated treating PQQ as a variable factor in the MaxPro design rather than a fixed background ingredient. Each step is recorded in the corresponding provenance manifest, allowing the recommendation to be re-traced from inputs to outputs.

**Figure 4.**
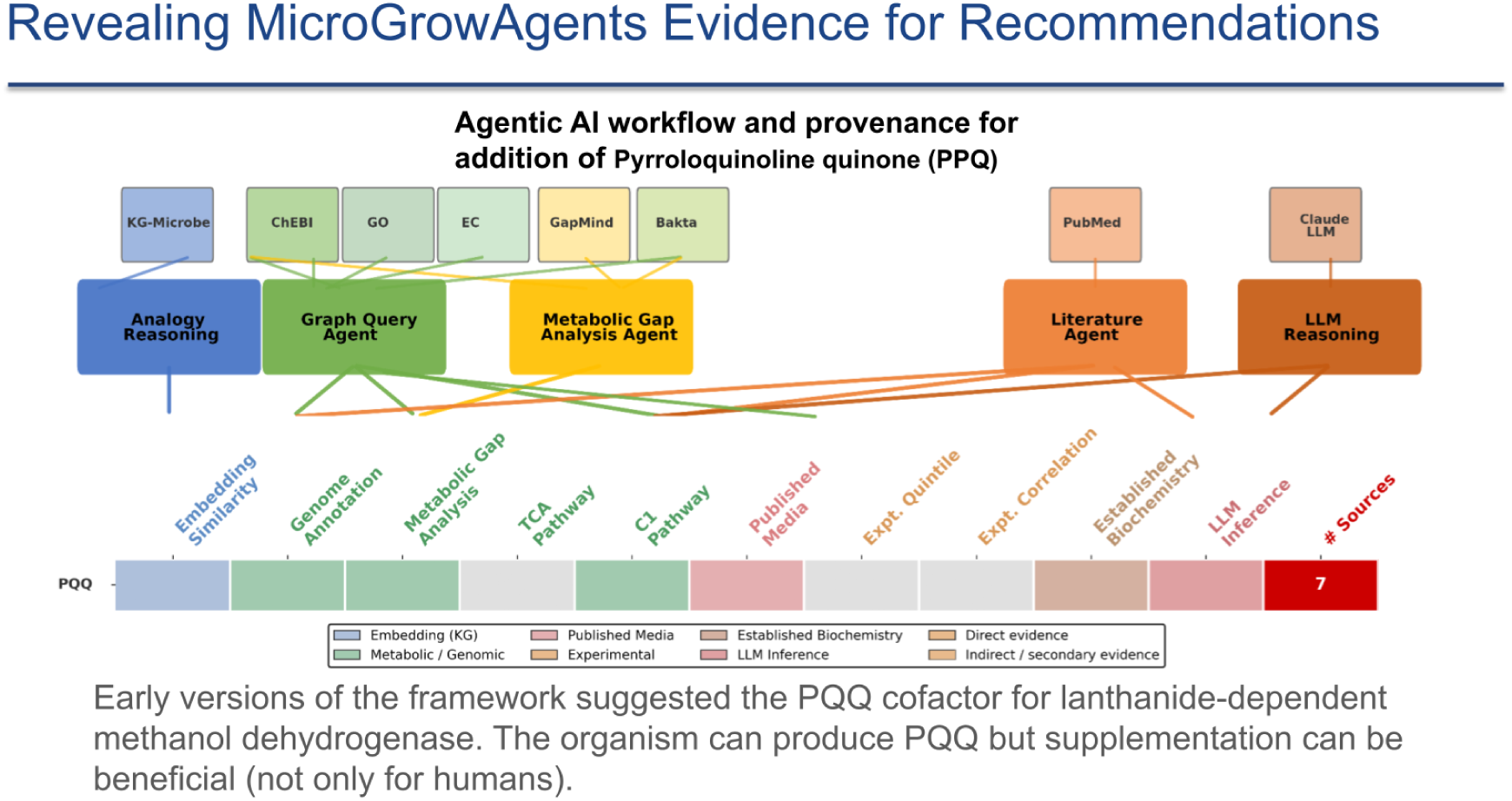
Agentic workflow and provenance trail by which MicroGrowAgents recommended pyrroloquinoline quinone (PQQ) as a variable factor in the M. extorquens AM1 MaxPro design. Five agents (Analogy Reasoning, Graph Query, Metabolic Gap Analysis, Literature, LLM Reasoning) contribute evidence from distinct data sources (KG-Microbe, ChEBI, GO, GapMind, Bakta, PubMed). The bottom strip lists per-agent evidence categories that feed the final aggregated recommendation: PQQ supplementation is beneficial despite AM1’s endogenous PQQ biosynthesis capability, motivating its inclusion as a designed factor rather than a fixed ingredient. Caveat (Supplementary Table S3): in the recorded design session the Graph Query agent’s KG response was a mock; the load-bearing evidence in the final aggregation came from the curated DOI-linked publication corpus, the AM1 Bakta annotations, and the curated FACTS sheet for AM1.

Response surface modeling using radial basis function (RBF) interpolation [regis2007rbf] enables identification of optimal nutrient combinations from experimental growth data by constructing smooth surfaces mapping nutrient concentrations to growth rates. Pairwise response surfaces across the six designed factors (Total_Phosphate, (NH₄)₂SO₄, CoCl₂, Succinate, Methanol, PQQ) were generated for visual inspection of ingredient interactions and used to guide downstream multi-objective Pareto analysis (next subsection).

### Round-2 Experimental Validation

The MaxPro design was executed in a Round-2 cultivation campaign of *M. extorquens* AM1 comprising 70 unique conditions (an updated working set from the 69-condition Round-1 design plus refinement candidates), each replicated four times across plates with randomized well positions and shared control wells. Cultures were inoculated into the designed media and tracked through three plate-reader timepoints; the t2 endpoint was used for primary analysis after abiotic-correction diagnostics indicated that t2 captured biomass signals before precipitation artifacts began to dominate at t3. Two response variables were recorded at each well: absorbance at 740 nm (cell-density proxy for biomass) and residual neodymium (Nd) concentration in spent medium quantified by arsenazo III colorimetric assay (with lower remaining Nd indicating higher apparent residual-Nd depletion). The growth curves on Biolog Odin were used to derive area under the curve (AUC), maximum specific growth rate (μ_max) and maximum absorbance at 740 nm. Relative to the standard MP base medium for this organism — the unsupplemented starting-point recipe carried as ctrl_media on every plate — the MC-stable composite candidate MPOB_058 increased integrated biomass by 79% (AUC at 740 nm: 5.12 ± 0.13 vs 2.86 ± 0.24), peak biomass by 97% (0.152 ± 0.007 vs 0.077 ± 0.007), and maximum specific growth rate by 46% (μ_max: 0.247 ± 0.009 vs 0.169 ± 0.015 h⁻¹; n = 4 vs 16 replicates; Gompertz fits, R² > 0.99). Because these kinetic metrics integrate the full 48 h time-course, they are larger than the single t2 endpoint OD600 difference used for the two-objective Pareto ranking (Table 1). The Biolog Odin growth curves and pairwise response surfaces for these conditions are shown in Figure 5, with an orthogonal 48-well shaken-microplate validation of the top five candidates in Figure 7. The 79%/97%/46% improvement above is measured against the on-plate MP base control rather than against a literature-optimized AM1 medium tuned for lanthanide-depletion growth; comparison to such a stronger comparator awaits a future round.

**Figure 5.**
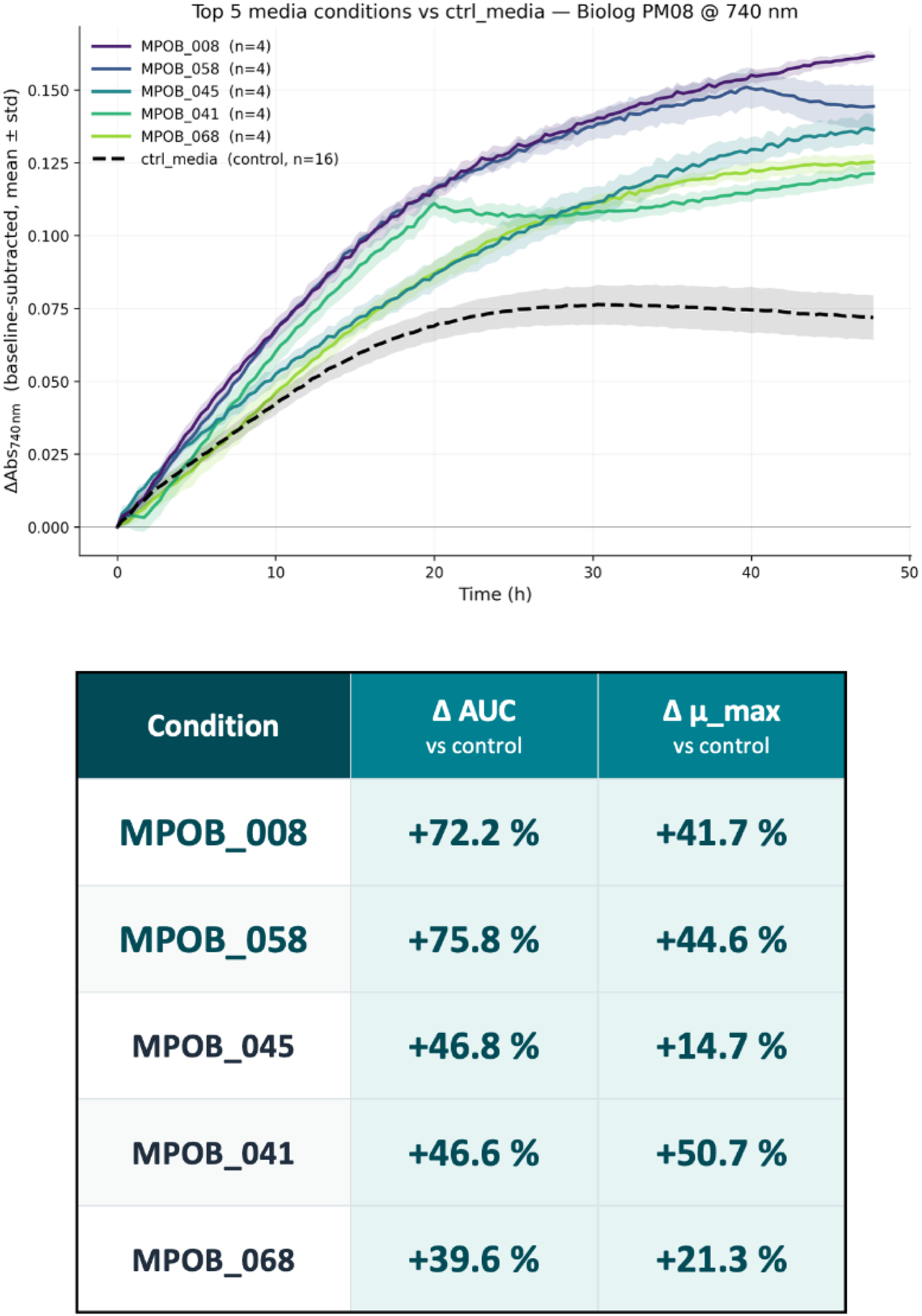
Biolog PM08 cultivation of the top-five MaxPro media designs for Methylorubrum extorquens AM1 (740 nm turbidity, n = 4 replicates per design; n = 16 for the ctrl_media baseline). Mean ± SD over the 48-hour cultivation window. The inset table reports Δ AUC and Δ μ_max for each design relative to the on-plate control.

**Figure 6.**
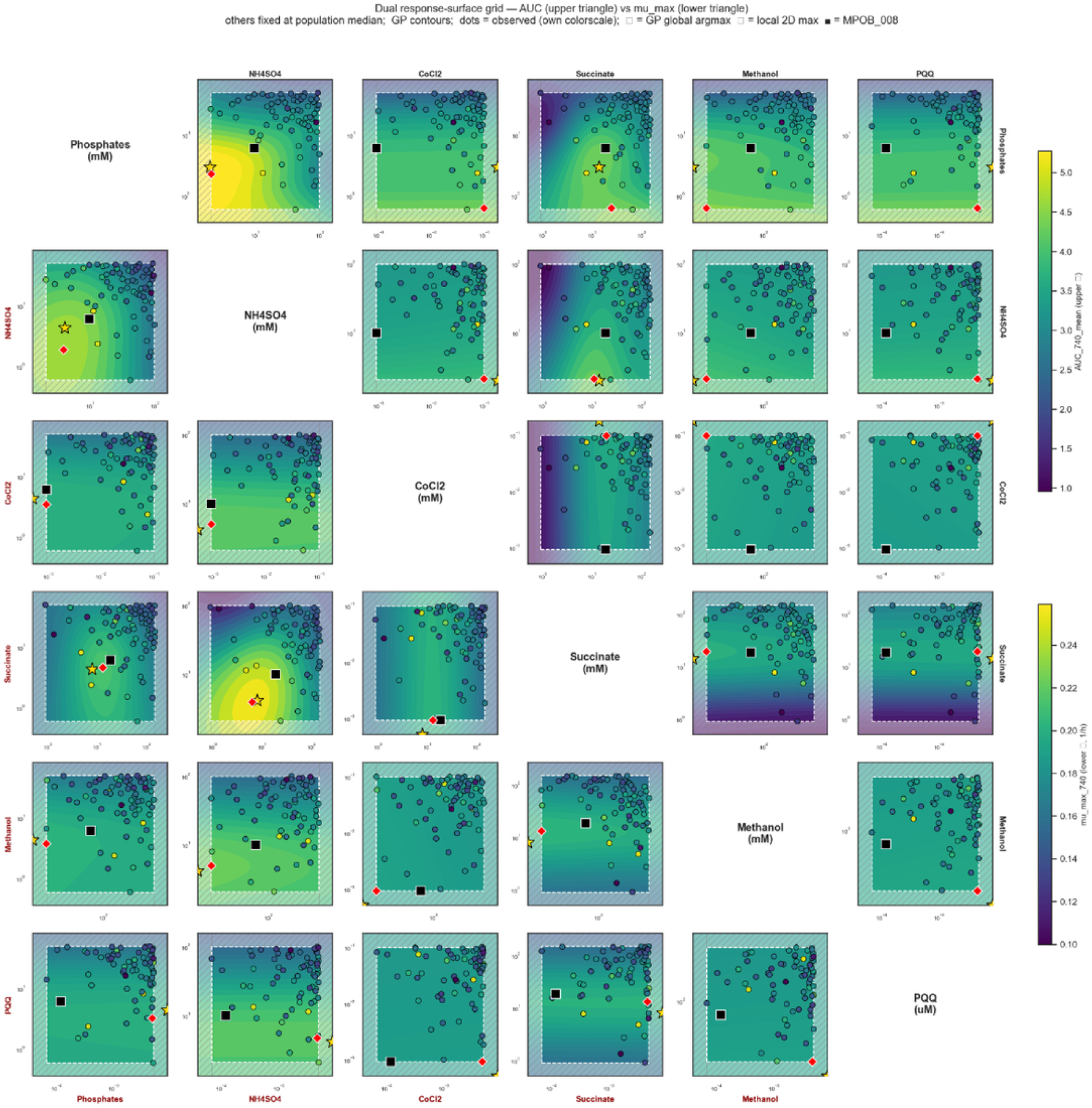
Pairwise response-surface grid for the six designed factors (Total_Phosphate, (NH₄)₂SO₄, CoCl₂, Succinate, Methanol, PQQ) in Round-2 cultivation of Methylorubrum extorquens AM1. Upper triangle: integrated biomass (AUC at 740 nm); lower triangle: maximum specific growth rate (μ_max). Each off-diagonal panel projects the 70 Round-2 conditions onto a pair of factors with the remaining four held at their population medians; GP-derived contours overlay the observed values (dots, own colourscale). Yellow stars mark the per-panel GP global argmax; black squares mark MPOB_008. Factor pairs with the steepest gradients identify the combinations that drive Round-2 medium performance.

**Figure 7.**
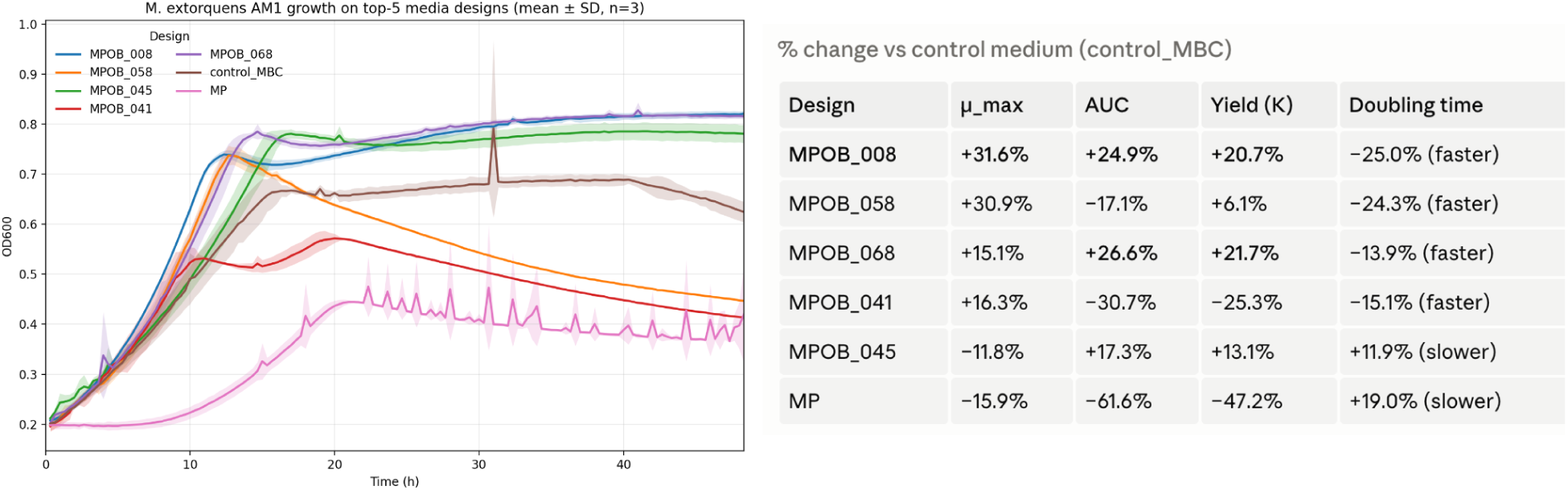
Orthogonal 48-well shaken-microplate validation of the top-five MaxPro media designs for Methylorubrum extorquens AM1 (n = 3 replicates per design), plus control_MBC and a standard MP control. OD600 mean ± SD over the 48-hour cultivation window. The inset table reports percent change versus control_MBC in μ_max, AUC, yield (K), and doubling time. Panel 7 confirms the Biolog Odin ranking (Figure 5) under an orthogonal cultivation geometry and demonstrates that the Round-2 Pareto-optimal media reproduce outside the Biolog format.

**Table 1.** Monte-Carlo Pareto frontier membership for Round-2 conditions surviving resampling (1000 iterations, endpoint t2). Objectives: maximize OD600 (biomass at 740 nm) and minimize Nd_uM remaining (apparent residual-Nd depletion). Values are means ± replicate standard deviation. “Det. Pareto” indicates whether the condition lies on the deterministic Pareto front. Stability tiers: stable (MC freq ≥ 0.8), borderline (0.5 ≤ freq < 0.8), rare (0.1 ≤ freq < 0.5).

| Condition | OD600 $\pm \sigma$ | Nd remaining ($\mu\text{M}$) $\pm \sigma$ | MC freq | Det. Pareto | Tier |
| --- | --- | --- | --- | --- | --- |
| MPOB_058 | 0.241 ± 0.003 | 2.58 ± 0.84 | 0.922 | ✓ | stable |
| MPOB_008 | 0.231 ± 0.007 | 2.07 ± 0.46 | 0.677 | ✓ | borderline |
| MPOB_019 | 0.208 ± 0.007 | 1.70 ± 0.39 | 0.514 | ✓ | borderline |
| MPOB_002 | 0.200 ± 0.016 | 2.74 ± 1.37 | 0.226 | — | rare |
| ctrl_media | 0.205 ± 0.010 | 2.52 ± 0.70 | 0.142 | — | rare |
| MPOB_041 | 0.226 ± 0.006 | 3.43 ± 0.90 | 0.100 | — | rare |

Identifying the best medium under two simultaneous and partly competing objectives is a multi-objective optimization problem: maximize biomass and minimize Nd remaining in spent media. We applied the Pareto front concept: the set of conditions for which no other condition simultaneously matches or exceeds performance on both objectives. Computed from replicate-mean endpoint values alone, the deterministic Pareto front comprised three conditions (Supplementary Table S1): MPOB_058 (OD600 0.241, Nd remaining 2.58 µM), MPOB_008 (0.231 / 2.07 µM), and MPOB_019 (0.208 / 1.70 µM). These three conditions trade off biomass against apparent residual-Nd depletion: MPOB_058 maximizes growth while leaving more Nd in solution, whereas MPOB_019 produces lower biomass but achieves the deepest lanthanide depletion.

Deterministic Pareto membership computed from replicate means alone can be brittle: a condition can appear on the front simply because of favorable replicate noise on a single axis. To quantify robustness under replicate-level uncertainty, we ran a Monte-Carlo Pareto frontier analysis (1000 iterations). For each iteration, every condition’s two coordinates were independently resampled from normal distributions parameterized by the observed replicate mean and standard deviation; the Pareto front was recomputed; and each condition’s frequency of appearance across the 1000 fronts was recorded (Figure 8). The histogram of frequencies shows a strongly bimodal distribution: most conditions never appear on the front (frequency < 0.1, grey), a small tail of rare appearers (0.1–0.5, light orange) reflects candidates whose front membership depends on lucky replicate variance, and only a handful of conditions appear consistently. Using stability tiers defined a priori (stable ≥ 0.8, borderline 0.5–0.8, rare 0.1–0.5, off-frontier < 0.1), MPOB_058 emerged as the sole stable winner (frequency 0.922), with MPOB_008 (0.677) and MPOB_019 (0.514) classified as borderline (Table 1). All three deterministic Pareto winners survived Monte Carlo (MC) resampling above the rare-appearer threshold, so the deterministic front is not an artifact of noise. However, the wide separation in MC frequency among the three winners — 0.922 vs. 0.677 vs. 0.514 — places confidence in MPOB_058 well above the other two.

**Figure 8.**
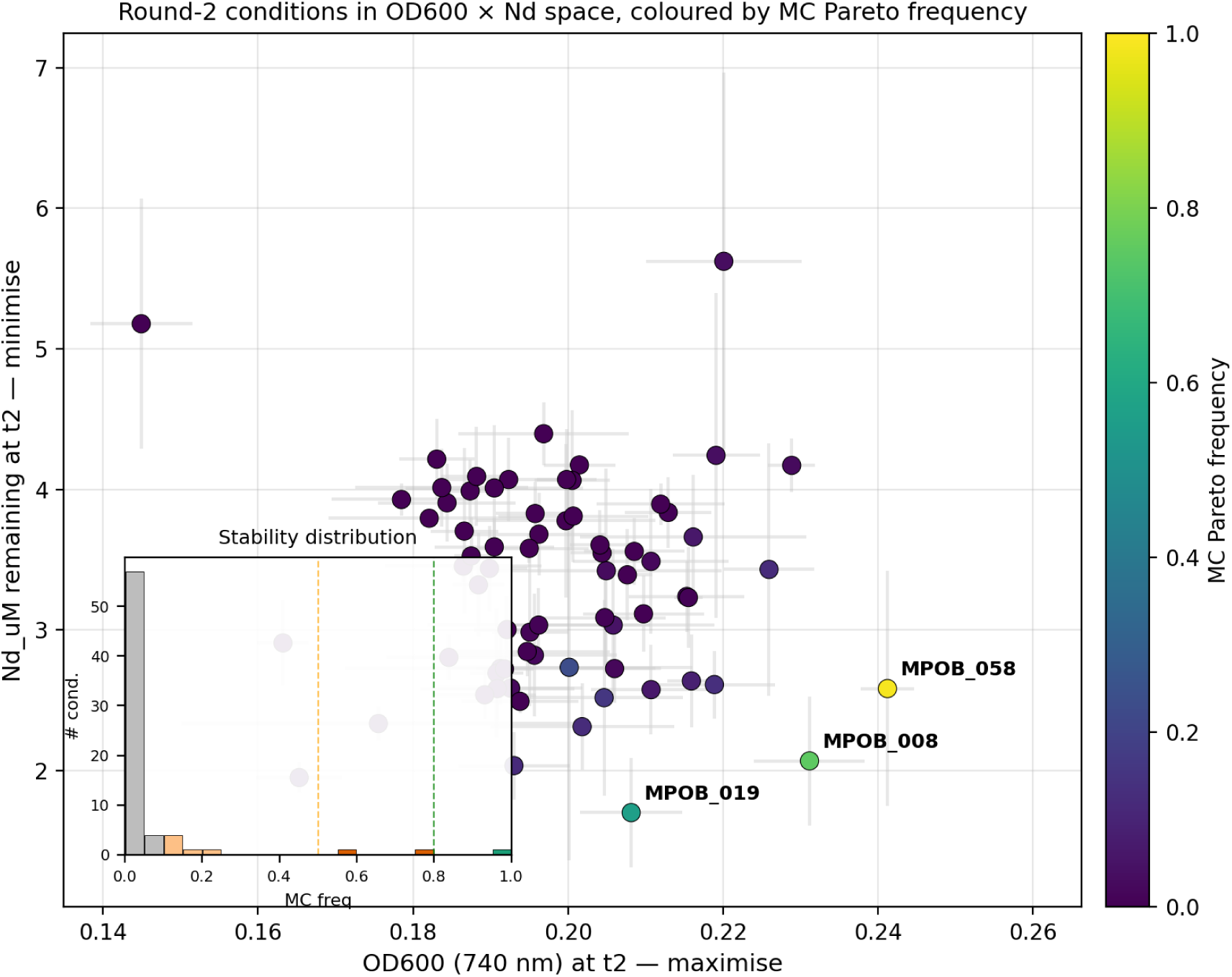
Monte-Carlo Pareto frontier. analysis of Round-2 M. extorquens AM1 cultivation (1000 iterations, t2 endpoint). Main panel: OD600 × Nd_uM remaining scatter with replicate σ as error bars, color-coded by MC Pareto-membership frequency on a viridis scale; stable and borderline winners (MPOB_058, MPOB_008, MPOB_019) are labelled. Inset: histogram of Pareto-membership frequencies across all 70 conditions, with bars colored by stability tier (green = stable ≥ 0.8, orange = borderline 0.5–0.8, light orange = rare 0.1–0.5, grey = off-frontier < 0.1); dashed vertical lines mark the stable (0.8) and borderline (0.5) thresholds. Objectives: maximize OD600 (740 nm biomass) and minimize Nd remaining (= maximize uptake).

Visualizing the OD600 × Nd_uM projection (Figure 8) with replicate σ as error bars and viridis coloring by MC frequency clarifies the structure of the design space: MPOB_058 occupies a corner combining high biomass with moderate Nd uptake; MPOB_019 trades biomass for the lowest Nd remaining; and the dense cloud of off-frontier conditions (grey) clusters at lower OD600 and intermediate Nd, separated from the winners by a clear gap. The inset histogram in Figure 8 summarizes the same MC frequency distribution across all 70 conditions, with the four stability tiers shaded by color. The pairwise response surface for the methanol × PQQ factor pair (Supplementary Figure S1), the two factors most directly relevant to lanthanide-independent methanol dehydrogenase activity, shows a smooth monotonic ridge of high OD600 at intermediate-to-high methanol and elevated PQQ, consistent with a role for PQQ-dependent methanol dehydrogenase under the surveyed lanthanide-depletion conditions. Both the Ca²⁺-dependent MxaF and the lanthanide-dependent XoxF methanol dehydrogenases require PQQ, and strain AM1 encodes the full PQQ biosynthetic operon, so supplemental PQQ is best read as a designed probe rather than a demonstrated rate limitation.

A paired-control biology analysis at the t2 endpoint refines the interpretation: MPOB_008 borderline-passes all three independent filters (MC stability, t2 biology signal −0.0197, near-clean abiotic baseline) and is the strongest biological-uptake candidate, while MPOB_058 is MC-stable and shows a t2 biology signal (−0.0095) but is chemistry-confounded (abiotic drift ≈4× any other Pareto winner). The Round-2 results therefore yield a prioritized anchor set — MPOB_058 as the composite stability winner suitable for paired ICP-MS to disentangle real biology from real chemistry, MPOB_008 as the cleanest biological-uptake candidate for replication-heavy follow-up, and MPOB_019 as a borderline-stable candidate whose ranking needs confirmation — together with a quantified estimate of where in the six-dimensional parameter space the next DBTL round should concentrate experimental effort.

**Supplementary Figure S1.**
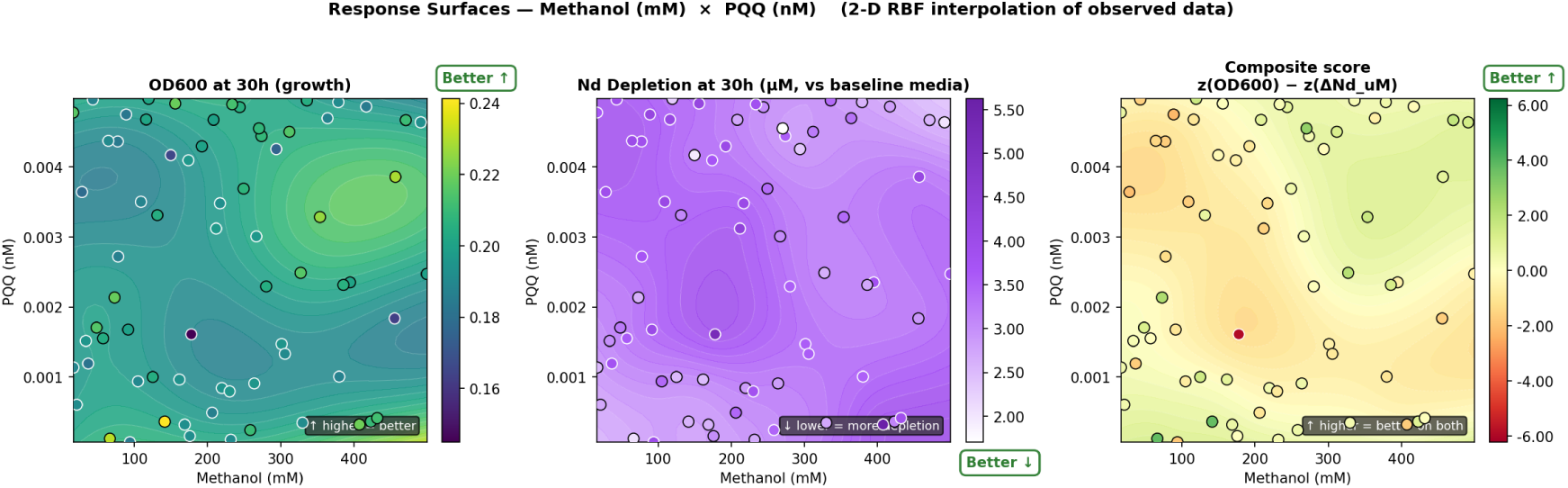
Pairwise response surface for the methanol × PQQ factor pair in Round-2 M. extorquens AM1 cultivation, fit by radial basis function interpolation over the t2 OD600 endpoint with the remaining four MaxPro factors held at their design medians. The high-OD600 ridge at elevated PQQ concentration is consistent with, and hypothesis-generating for, a role of PQQ-dependent methanol dehydrogenase under the surveyed lanthanide-depletion conditions (both MxaF and XoxF require PQQ; AM1 encodes the full pqq operon), and matches the agentic recommendation that motivated PQQ’s inclusion as a designed factor (Figure 4).

**Supplementary Table S1.** Deterministic two-objective Pareto-optimal conditions identified from Round-2 *M. extorquens* AM1 cultivation, computed from replicate-mean endpoint measurements. All three deterministic winners survive Monte-Carlo resampling above the rare-appearer threshold (Table 1).

| Condition | OD600 (740 nm) | Nd remaining ($\mu\text{M}$) |
| --- | --- | --- |
| MPOB_058 | 0.241 | 2.58 |
| MPOB_008 | 0.231 | 2.07 |

| Condition | OD600 (740 nm) | Nd remaining<br>($\mu\text{M}$) |
| --- | --- | --- |
| MPOB_019 | 0.208 | 1.70 |

### Held-out Predictive Validation of the Design Factors

Because the six designed factors — not the agentic interpretation layer — set the experimental search space, we asked whether those factors carry learnable signal about growth, and how that signal compares across modeling strategies. Using leave-one-condition-out cross-validation over the 70 Round-2 conditions, we predicted each held-out condition’s integrated biomass (AUC at 740 nm) from its six factor concentrations. Gradient boosting, random forests, and Gaussian-process regression all substantially outperformed a naive mean predictor (R² = 0.47–0.58 vs 0.00 for the naive baseline) and recovered three to five of the five highest-growth conditions (Precision@5 0.60–1.00; Supplementary Table S2; benchmark code at manuscript/MicroGrowAgents_v8/benchmark/run_dry_benchmark.py). An independent Automated Recommendation Tool (ART) cross-validation on the replicate-level data reached R² ≈ 0.92. By contrast, a constraint-based flux-balance baseline could not serve as a quantitative predictor here. The available strain AM1 genome-scale reconstruction is an un-gapfilled draft: it lacks a connected methanol-oxidation pathway (methanol enters the cytosol with no downstream fate), does not represent the lanthanide/PQQ-dependent methanol-dehydrogenase chemistry at all, and does not support growth on a defined methanol minimal medium. Moreover, across the surveyed ranges the designed factors are saturating (methanol, succinate) or above biomass demand (phosphate, sulfate, cobalt), so even a fully curated stoichiometric model would not differentiate the conditions without an added uptake-kinetics layer. The structure that ranks candidate media in this design space is therefore statistical-empirical rather than recoverable from constraint-based stoichiometry alone, and curating a methylotrophic genome-scale model for quantitative growth prediction remains future work. We did not benchmark across rounds: Round-1 and Round-2 used different measurement modalities (600 nm single-channel absorbance versus 740 nm Biolog turbidity), the source of the near-zero cross-round rank correlation, so a held-out comparison within Round-2 is the defensible test. This benchmark establishes that the design factors are genuinely informative about Round-2 growth and that data-driven surrogates, rather than mechanistic FBA, capture the structure that prioritizes candidate media. We emphasize that this benchmark validates the factor space — the six designed variables and their concentration ranges — not the agentic interpretation layer that selected them. The agentic contribution is tested separately via a retrospective comparison against a deterministic literature/MediaDive baseline.

**Supplementary Table S2.**
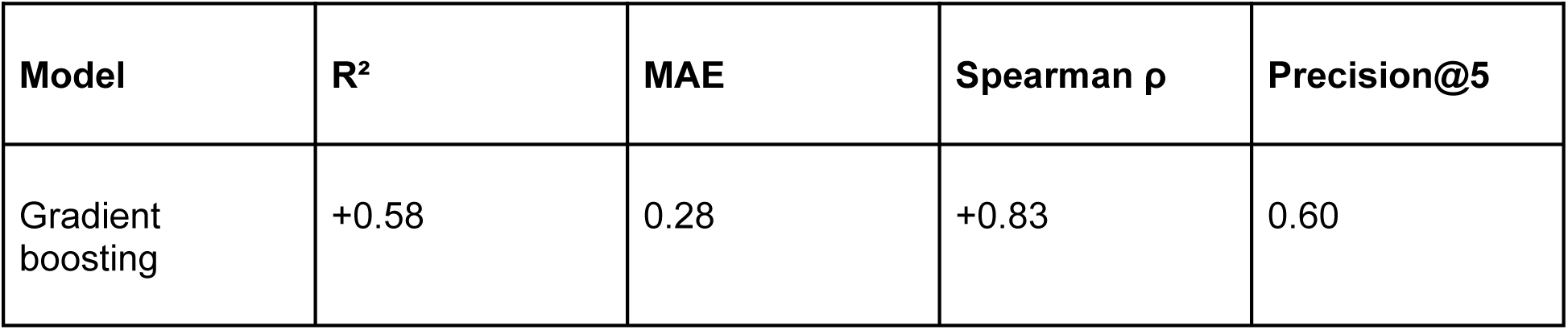
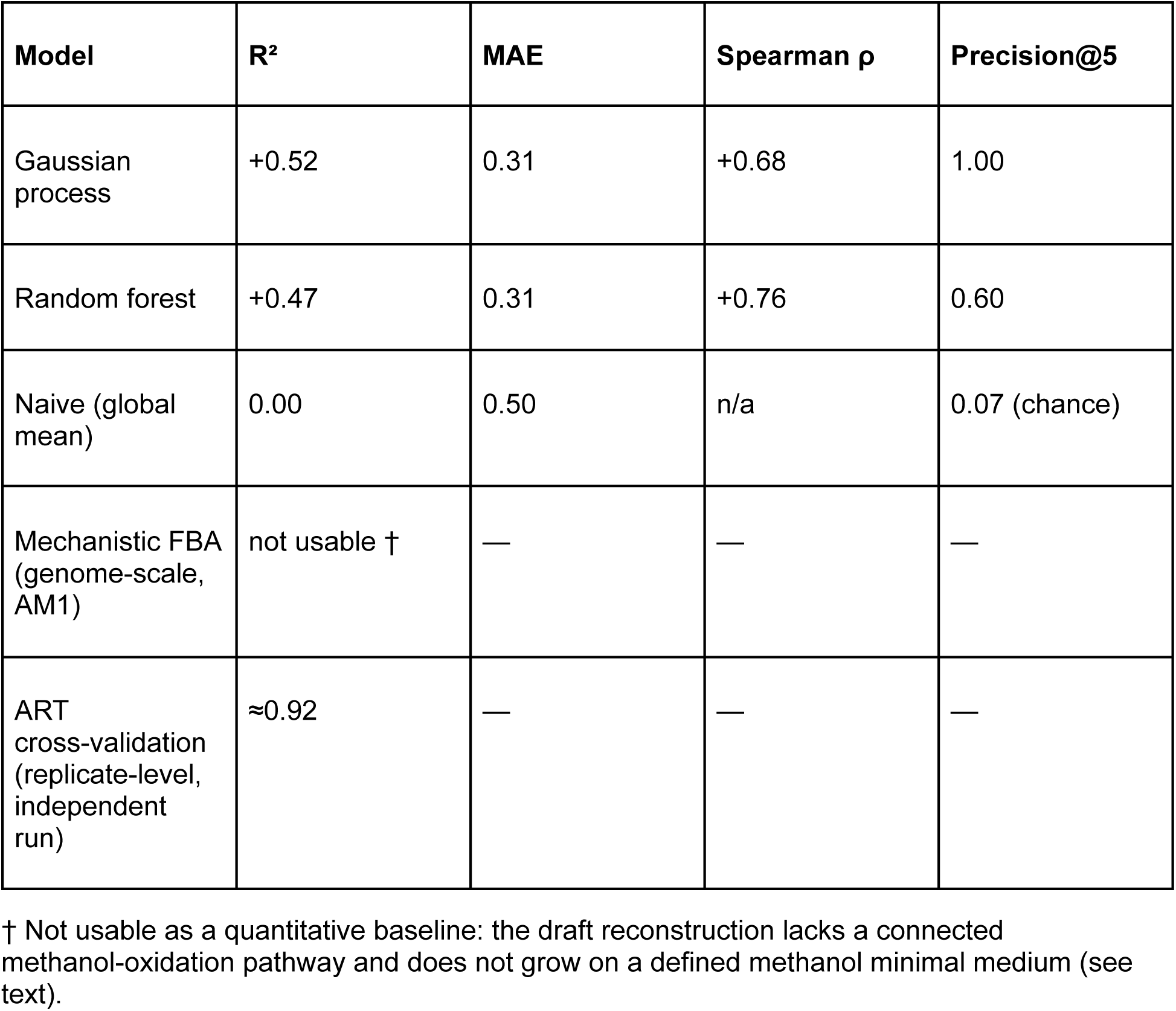
Held-out (leave-one-condition-out) prediction of Round-2 integrated biomass (AUC at 740 nm) from the six designed factors, over 70 conditions. Higher R², Spearman ρ, and Precision@5 (recovery of the true top-5 conditions) and lower MAE are better.

| Model | $R^2$ | MAE | Spearman $\rho$ | Precision@5 |
| --- | --- | --- | --- | --- |
| Gradient boosting | +0.58 | 0.28 | +0.83 | 0.60 |

| Model | R<sup>2</sup> | MAE | Spearman $\rho$ | Precision@5 |
| --- | --- | --- | --- | --- |
| Gaussian process | +0.52 | 0.31 | +0.68 | 1.00 |
| Random forest | +0.47 | 0.31 | +0.76 | 0.60 |
| Naive (global mean) | 0.00 | 0.50 | n/a | 0.07 (chance) |
| Mechanistic FBA (genome-scale, AM1) | not usable † | — | — | — |
| ART cross-validation (replicate-level, independent run) | ≈0.92 | — | — | — |

† Not usable as a quantitative baseline: the draft reconstruction lacks a connected methanol-oxidation pathway and does not grow on a defined methanol minimal medium (see text).

### Retrospective comparison to a deterministic literature/MediaDive baseline

The held-out predictive benchmark above validates the six designed factors as a search space, but does not address the related question reviewers and editors are likely to ask: would a deterministic non-agentic pipeline have selected the same six factors from the same input data, or did the agentic interpretation layer actually change which variables entered the design? To probe this, we constructed a deterministic literature/MediaDive baseline (“Det-MD”) whose factor selection uses only public MediaDive media records and the same MP-base ingredient list available to the agentic pipeline, with no LLM agent in the loop, and compared its choices to those that MicroGrowAgents made.

We ran two Det-MD arms. Det-MD-A filters MediaDive to media that the underlying knowledge graph documents as supporting a methylotroph organism (15 genus prefixes: Methylorubrum, Methylobacterium, Methylobacillus, Methylophilus, Methylopila, Methylocystis, Methylocella, Methylomonas, Methylococcus, Methylosinus, Methylocaldum, Methyloferula, Methylobacter, Methylomicrobium, Methylotenera; 3,116 organism records, 37 distinct media). Det-MD-B widens the filter to media that contain any C1 carbon source (methanol, formate, methylamine, or formaldehyde; 301 media) as a robustness check. Both arms exclude the 14 MP-base infrastructure ingredients (buffers, trace metals, gelling agents) by the same pattern list used by the agentic pipeline, then rank the remaining ingredients by the number of distinct media they appear in, and take the top six as the Det-MD factor set. Concentration ranges are reported as g/L median/IQR within the dominant unit because MediaDive concentrations live in heterogeneous units (g/L, ml, %(v/v), mM) and the underlying ingredient catalog’s molecular weights are mostly a sentinel 100.0 g/mol, making mmol/L conversion unreliable across the corpus. Both arms reuse the same MaxPro+OptBlock generator and the same Monte-Carlo Pareto routine that the agentic arm used; only the input factor table differs. The Det-MD logic (genus list, fixed-ingredient patterns, top-N cutoff) was committed and tagged (ablation-prereg-v1) before running against Round-2 outcomes; the commit hash and clean-worktree flag are recorded in outputs/ablation_det_md/RESULTS.json.

Det-MD-A recovered four of the six MGA-chosen factors — Total_Phosphate, (NH₄)₂SO₄, CoCl₂, and methanol — purely from organism-frequency ranking, with no agentic step. The Jaccard agreement on the factor set is 0.50. The two MGA-only factors are succinate and PQQ (Supplementary Table S4, Supplementary Figure S2). Det-MD-A’s two non-MGA selections — sodium chloride and glucose — illustrate where frequency-only ranking misfires for AM1: NaCl is high-frequency in MediaDive’s methylotroph subset largely because the corpus includes marine and brackish-water isolates whose biology does not transfer to AM1, and glucose is a generic-microbiology default that AM1 (an obligate methylotroph on C1 substrates) cannot use as a primary carbon source. Det-MD-B widens the medium pool to 301 C1-substrate media and confirms the qualitative pattern: the same three MGA factors remain shared (phosphate, ammonium sulfate, methanol), with CoCl₂ now also flagged as MGA-only because the broader C1 set dilutes the cobalt-cofactor signal that the methylotroph-specific filter preserves.

The interpretation we draw is narrower than the agentic-discovery claim a high-impact reading of the manuscript might invite. Four of MicroGrowAgents’s six factor choices are recoverable from MediaDive frequency alone, so the agentic contribution is not “factor discovery” in those cases. The two factors that survive only the agentic pipeline are precisely those whose justifications live outside MediaDive: PQQ as a curated-FACTS- and Bakta-annotated cofactor of methanol dehydrogenase (Figure 4), and succinate as a TCA-cycle replenishment carbon source motivated by the AM1 curated FACTS sheet rather than by MediaDive’s media catalog. The agentic layer’s load-bearing contributions in this case study are therefore (i) selecting two factors that the deterministic baseline misses because the supporting evidence is in literature, genome, and curated knowledge rather than in a media catalog; (ii) avoiding two factors that the deterministic baseline picks but that biology rules out for AM1; and (iii) the auditable evidence trail that lets a reviewer re-derive each decision. We treat this as the more defensible scope of the contribution and adopt it as the framing for the Discussion (“Deterministic core, agentic edges”), the Bigger Picture, and the title.

**Supplementary Table S4.** Det-MD baseline factor selection compared to MicroGrowAgents’s agentic factor selection for the Methylorubrum extorquens AM1 case study. Det-MD-A: methylotroph organism filter (3,116 organism records → 37 distinct media). Det-MD-B: C1 carbon source filter (301 media). Both arms exclude the 14 MP-base infrastructure ingredients and rank the remaining ingredients by number of distinct media of occurrence; concentration ranges are g/L within the dominant unit because MediaDive’s ingredient catalog lacks reliable molecular-weight values for mmol/L conversion. The Det-MD code is pre-registered under git tag ablation-prereg-v1 in the source repository; RESULTS.json records the commit hash captured at run time.

| MGA factor | In Det-MD-A? | Det-MD-A range (g/L, p25–p75) | In Det-MD-B? | Det-MD-B range (g/L, p25–p75) | Comment |
| --- | --- | --- | --- | --- | --- |
| Total_Phosphate | yes | 0.54–2.97 | yes | (see RESULTS.json) | recovered by frequency |
| (NH<sub>4</sub>)<sub>2</sub>SO<sub>4</sub> | yes | 0.50–2.00 | yes (via NH<sub>4</sub>Cl bucket) | (see RESULTS.json) | recovered by frequency |
| CoCl<sub>2</sub> | yes | 0.12–0.20 | no | — | recovered only with organism filter |
| Methanol | yes | 4.79–7.92 | yes | 4.79–9.48 | recovered by both filters |
| **Succinate** | **no** | — | **no** | — | **MGA-only; curated FACTS evidence** |
| **PQQ** | **no** | — | **no** | — | **MGA-only; literature + Bakta evidence** |
| Factor Jaccard vs MGA |  | 0.50 |  | 0.33 |  |
| Det-MD factors not in MGA |  | NaCl, Glucose |  | NaCl, Na<sub>2</sub>S, NaHCO<sub>3</sub> | frequency artefacts |

**Supplementary Figure S2.**
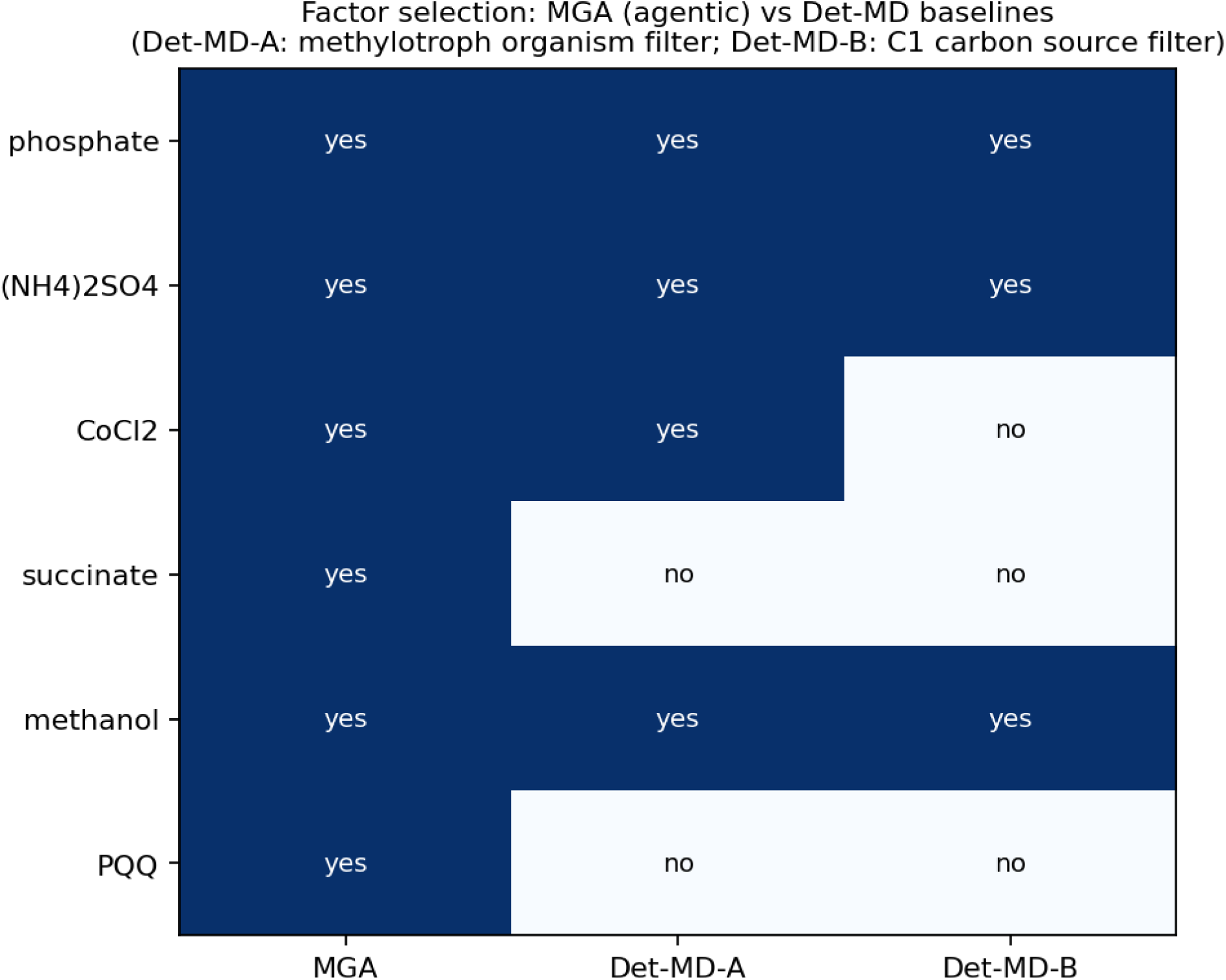
Factor-selection comparison: MGA (agentic, all six yes) vs Det-MD-A (methylotroph organism filter; selects phosphate, (NH₄)₂SO₄, CoCl₂, methanol; misses succinate, PQQ) vs Det-MD-B (C1 carbon source filter; selects phosphate, (NH₄)₂SO₄ via the NH₄Cl bucket, methanol; misses CoCl₂, succinate, PQQ). The binary selection table makes the load-bearing agentic contributions — PQQ and succinate — directly visible.

### Tasks Where the Agentic Framework Contributed

Across the v10 design and its two cultivation rounds, several pipeline decisions were shaped by the agentic framework — its curated knowledge, evidence aggregation, and interpretation of tool output — rather than by a fixed algorithm alone (Supplementary Table S3). These span the full design–build–test–learn cycle: surfacing and promoting a candidate cofactor (PQQ) from a background ingredient to a designed factor, reconciling a contradictory auxotrophy call against curated genome facts, re-scoping factor ranges between rounds, resolving an instrumentation labeling mismatch, selecting the analysis endpoint, and converting the Round-2 Pareto analysis into a prioritized anchor set for Round-3. The design-stage decisions are captured in the documented session manifests (see Data Provenance and Reproducibility, below); the Round-2 analyses were run as deterministic scripts whose outputs were interpreted to seed the next round, so each recommendation can still be re-traced from inputs to outputs.

Agentic value is not confined to steps that lack a deterministic tool. Many of the tasks in Supplementary Table S3 invoke ordinary Python scripts and standard bioinformatics tools — COBRApy flux balance analysis, GapMind, RDKit fingerprinting, the MaxPro generator, the Monte-Carlo Pareto routine — yet the agentic framework still contributes three capabilities a bare script call does not. First, error tolerance: when a tool fails, returns an implausible value, or contradicts curated knowledge — as when GapMind reported a spurious cobalamin auxotrophy, or when arsenazo readings went negative under missing calibration — the agent detects the inconsistency and routes around it rather than propagating it downstream. Second, output interpretation: raw tool output (an FBA flux vector, an RBF response surface, a Monte-Carlo frequency histogram) is read in biological context to decide what it means for the next action, for example reading the methanol × PQQ response-surface ridge as consistent with elevated demand on PQQ-dependent methanol dehydrogenase under lanthanide depletion (AM1 encodes the complete PQQ biosynthetic operon, so this is hypothesis-generating rather than a demonstrated rate limitation). Third, adaptive reruns: the agent re-parameterizes and re-invokes the same tool to improve a result, as in the Round-1 → Round-2 factor-range re-design and the re-extraction of kinetic timepoints onto the Round-1 endpoint schema. In each case the underlying computation remains deterministic and reproducible (Discussion, “Deterministic core, agentic edges”); the agentic contribution is the surrounding judgement about when to trust, re-interpret, or repeat it.

**Supplementary Table S3.** Pipeline tasks across the M. extorquens AM1 case study where the agentic framework contributed, the responsible component, and the resulting decision or output. Contributions are of three kinds: evidence-aggregation and reconciliation decisions taken in the documented design sessions (rows 1–4, partly human-in-the-loop), curated organism knowledge that corrected automated annotation, and deterministic analysis scripts whose outputs were interpreted to drive the next design–build–test–learn round (rows 5–9). Several tasks wrap deterministic scripts or standard bioinformatics tools; the framework’s contribution lies in evidence aggregation, error tolerance, biological interpretation of tool output, and adaptive re-invocation, not in replacing the underlying computation.

| Task | Framework component / role | Result |
| --- | --- | --- |
| Cofactor requirement survey | Curated cofactor reference (ChEBI/KEGG/BRENDA) feeding CofactorMediaAgent | 44 essential cofactors classified into six functional categories; PQQ surfaced among methylotrophy cofactors |
| Promote PQQ to a designed factor | Design-session evidence-aggregation decision over curated PQQ literature and genome lanthanide/PQQ-operon evidence (LanthanideGenesAgent); the KG graph query returned mock data | PQQ moved from fixed background ingredient to a variable MaxPro factor (Figure 4); Round-2 response surface confirmed a methanol × PQQ growth ridge (Figure 6) |
| Cobalamin auxotrophy reconciliation | Curated AM1 FACTS sheet weighed against GapMind's | Spurious “incomplete B<sub>12</sub> pathway” call corrected; v10 |
|  | partial-pathway call (knowledge layer) | design varies CoCl<sub>2</sub> rather than substituting vitamin B<sub>12</sub> |
| Designed-factor selection and range setting | Human-in-the-loop selection (logged design-session UserInput) informed by genome and cofactor analysis | Six nutrients chosen as variable factors with literature/MediaDive-derived ranges; genome and curated-FACTS evidence set which cofactors were varied rather than held fixed |
| Round-1 → Round-2 factor-range re-design | Deterministic re-design over Round-1 response surfaces, interpreted by the author | Round-2 factor ranges re-scoped from Round-1 dose-response, heatmap, and PCA outputs |
| Well-label resolution audit | Manual/script label audit (plot_swapped_methanol_succinate.py) | Position-labeling mismatch corrected; _SWAPPED_LABELS_* pre-resolution snapshots retained for traceability |
| Analysis-endpoint selection | Deterministic abiotic-correction script (analyze_round2_abiotic_correction.py), interpreted | t2 (≈6 h) adopted as the canonical endpoint over t1/t3 |
| Round-2 Pareto interpretation and hand-off | Deterministic scripts (mc_pareto_round2.py; t2 paired-biology), interpreted | Prioritized Round-3 anchor set defined: MPOB_058 (composite stability), MPOB_008 (cleanest biological uptake), MPOB_019 (needs replication) |
| Cross-round reproducibility and platform-switch decision | Deterministic comparison script (compare_round1_vs_round2.py) plus author decision | Near-zero cross-round correlation attributed to the instrumentation switch; subsequent rounds anchored on Round-2 winners only |

### Data Provenance and Reproducibility

MicroGrowAgents records all agent actions in structured YAML manifests following a standardized schema. Each session directory contains three files: manifest.yaml (session metadata, goals, outcomes), actions.jsonl (action-by-action log in JSON Lines format), and summary.md (human-readable summary). The action log captures eight action types: read, search, analyze, modify, create, execute, query, and decision. A total of 13 provenance sessions span the development period from January 10 to February 18, 2026, documenting iterative refinement of media designs (v5 through v13), experimental design generation, and data analysis workflows. Each manifest records the Claude model version (claude-sonnet-4-5 or claude-opus-4-6), session duration, files modified, and per-session checksums of selected input data files. The v10 MaxPro design session (2026-02-05-mp-plus-v10) records the input ingredient file checksum, MaxPro algorithm parameters (n_candidates=100,000, criterion=‘maxpro’), and the output design matrix checksum.

The artifact cleanup policy retains final outputs and their generating scripts and removes intermediate files (temporary CSVs, draft plots) after validation. Manifests and human-readable summaries are tracked in Git; full per-action JSONL logs are retained locally and will be deposited with the archival release. Against the bbop-skills criteria for local-first agentic systems, the system passes 7 of 9 criteria (78%) in the February 2026 audit; the two open gaps are MCP-standardized tool exposure and full cryptographic hashing of all experimental input datasets, both of which are tracked items on the integration roadmap. We therefore describe the current provenance as strong but not yet complete.

**Supplementary Figure S3.**
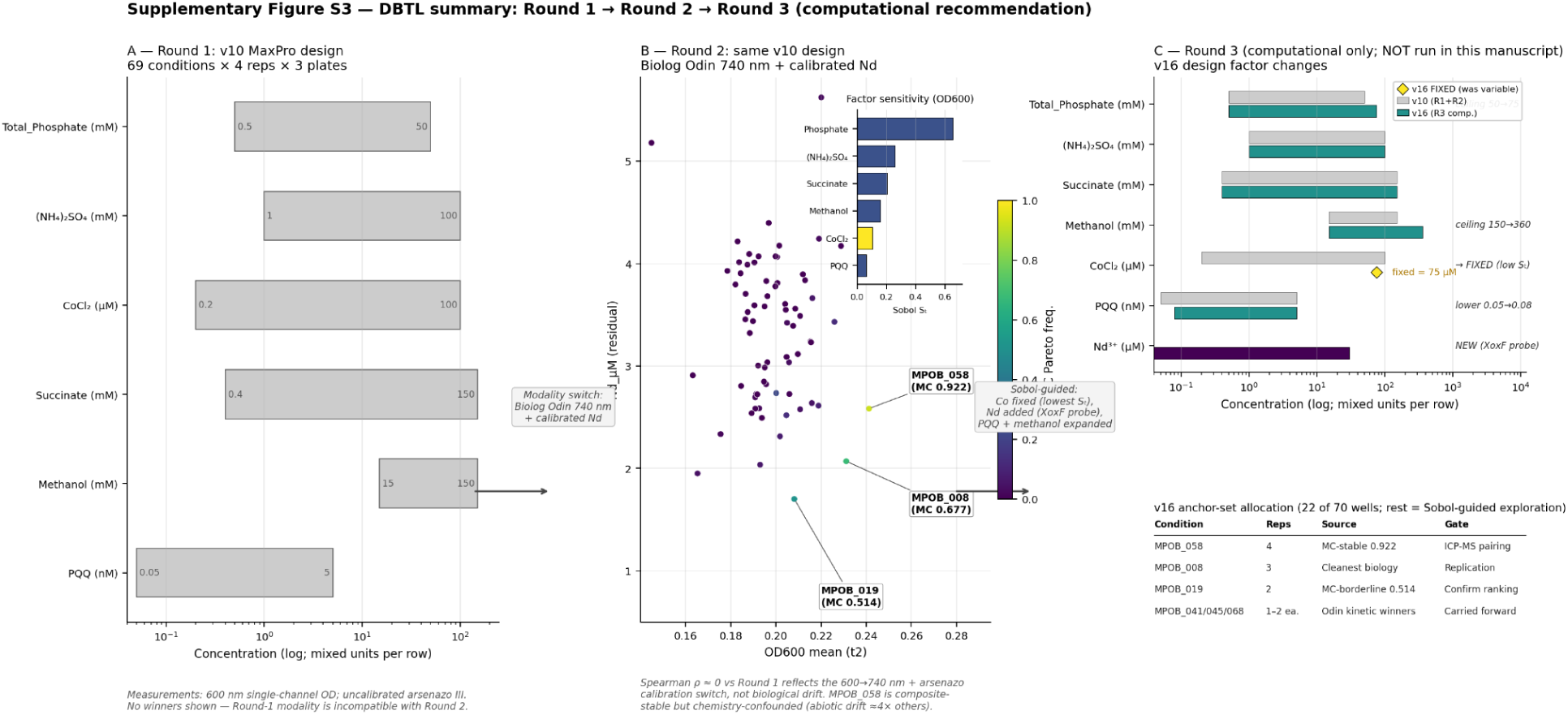
DBTL summary across Rounds 1–3 for Methylorubrum extorquens AM1. Panel A: Round 1 — v10 MaxPro design (69 conditions × 4 replicates × 3 plates) with 600 nm single-channel OD and uncalibrated arsenazo III; no winners are shown because the Round-1 measurement modality is incompatible with Round 2’s calibrated readout (Spearman ρ ≈ 0 across rounds, driven by the modality switch rather than biological drift). Panel B: Round 2 — same v10 design with calibrated Biolog Odin 740 nm turbidity and Miller-calibrated residual Nd. The three MC Pareto winners are labelled: MPOB_058 (stable, MC 0.922), MPOB_008 (borderline, 0.677), MPOB_019 (borderline, 0.514); the Sobol total-order sensitivity inset orders the six factors by their contribution to OD600 variance (Phosphate dominant, PQQ lowest). MPOB_058 is composite-stable but chemistry-confounded (abiotic drift ≈4× the other winners). Panel C: Round 3 — a computational-only v16 design recommendation (NOT run in this manuscript). Top sub-panel: per-factor concentration ranges, v10 (grey) versus v16 (teal), with range-refinement decisions annotated — Phosphate ceiling raised 50→75 mM, Methanol ceiling raised 150→360 mM, CoCl₂ moved from variable to fixed at 75 µM (lowest Sobol S□), PQQ lower-bound trace adjustment, and Nd³⁺ added as a NEW variable factor (0–30 µM) to probe XoxF-MDH lanthanide-dependent methylotrophy directly. Bottom sub-panel: the 22-well anchor-set allocation reusing the Round-2 winners under explicit follow-up gates (ICP-MS pairing on MPOB_058 to disambiguate chemistry vs biology; replication-heavy confirmation on MPOB_008; ranking confirmation on MPOB_019; carry-forward of the Odin kinetic winners MPOB_041, _045, _068). The remaining 48 wells are Sobol-guided exploration. Source data: outputs/round2_mc_pareto/ mc_pareto_membership.tsv, outputs/round1_vs_round2/REPRODUCIBILITY_REPORT.md, and data/designs/MP_latinhypercube/plate_designs_v16_maxprooptblock_long/. Generating script: scripts/figures/dbtl_summary_figure.py.

## Discussion

### Advantages of Agent-Based Architecture

The 29-agent, 58-skill organization lets each agent develop domain expertise in a specific subdomain (knowledge graph queries, metabolic modeling, experimental design), rather than diluting performance across a single monolithic model [majumdar2025]. Independent agents execute in parallel — KGReasoningAgent queries KG-Microbe while LiteratureAgent mines PubMed and GenomeFunctionAgent analyzes annotations — reducing total workflow time over sequential processing (specific speedup depends on workload and is not quantified here). Functional tiers also localize change: a ModelSEED schema update touches only the metabolic modeling tier instead of propagating across the codebase. The skill-based design admits new capabilities without refactoring: adding a MetaCyc pathway lookup means writing a new skill definition with its SPARQL or API query logic, mirroring successful modular bioinformatics frameworks [majumdar2025, ghafarollahi2024].

A second, less obvious advantage is that the agentic framework adds value even when the step it wraps is ultimately a deterministic Python script or a standard bioinformatics tool (COBRApy FBA, GapMind, RDKit, the MaxPro generator, the Monte-Carlo Pareto routine). Wrapping such a tool in an agentic, human-in-the-loop workflow does not change its computation, but it adds three capabilities a bare invocation lacks. The first is error tolerance: when a tool fails or returns an implausible or contradictory result, the agent detects the inconsistency and routes around it rather than propagating it — as when GapMind’s spurious cobalamin-auxotrophy call was reconciled against curated AM1 genome facts, or when uncalibrated arsenazo readings going negative flagged the need for a per-plate inverse calibration. The second is output interpretation: raw tool output is read in biological context to decide the next action, as when a methanol × PQQ response-surface ridge was read as hypothesis-generating for PQQ-dependent methanol-dehydrogenase activity under lanthanide depletion (not a demonstrated rate limitation, since AM1 encodes the full PQQ biosynthetic operon). The third is adaptive reruns: the agent re-parameterizes and re-invokes the same tool to improve a result, as in the Round-1 → Round-2 factor-range re-design. The deterministic core remains reproducible bit-for-bit (see “Deterministic core, agentic edges,” below); the agentic contribution is the surrounding judgement about when to trust, re-interpret, or repeat a tool’s output. Supplementary Table S3 enumerates the specific tasks in the M. extorquens AM1 case study where this kind of framework-level contribution shaped the outcome.

### Integration of Knowledge Graphs and Metabolic Models

The integration of knowledge graphs (KG-Microbe) with constraint-based metabolic models (COBRApy) addresses a fundamental challenge in media optimization: hypothesis generation. Traditional statistical optimization methods (DoE, Bayesian optimization) explore parameter spaces without biological context, treating nutrient concentrations as independent variables [joseph2025bayesian]. However, microbial growth depends on intricate metabolic networks where nutrient requirements exhibit complex dependencies. For example, cobalamin biosynthesis requires cobalt, but the specific requirement depends on the presence or absence of alternative B₁₂-independent metabolic pathways [seif2020]. Knowledge graphs provide this contextual information through structured relationships: KG-Microbe encodes organism-pathway-enzyme-cofactor relationships spanning 1.5M nodes and 5.1M edges [price2025kgmicrobe].

Genome-scale pathway-completeness and auxotrophy analysis enables qualitative hypothesis generation before experimental execution: for a given organism it flags which biomass precursors the genome can or cannot synthesize and suggests supplementation strategies [orth2010]. We did not apply quantitative flux-balance growth-rate prediction in this study — the available AM1 genome-scale reconstructions do not support methylotrophic growth (see Results) — so the metabolic-modeling layer is used here for qualitative gap reasoning rather than growth ranking. This pathway-completeness reasoning helps distinguish nutrients the organism must be supplied from those it can synthesize, informing which ingredients to hold fixed versus vary. In the *M. extorquens* AM1 case study, genome-scale pathway-gap analysis initially returned a partial-biosynthesis call for cobalamin (12 of 17 enzymes detected); the curated AM1 FACTS sheet established that AM1 encodes the complete de novo pathway, the apparent gap being an annotation artefact, so cobalt (CoCl₂) — the metal centre that pathway requires — was included as a variable ingredient rather than supplementing B₁₂. This illustrates curated knowledge correcting an automated annotation gap. Without this metabolic context, a purely statistical optimization might waste experimental budget exploring irrelevant nutrient combinations.

The integration of these complementary approaches—KG-Microbe for empirical knowledge, ModelSEED for biochemical reaction networks, and pathway-gap analysis for mechanistic feasibility checks—creates a multi-scale framework spanning molecular interactions to organism-level phenotypes. This mirrors successful systems biology approaches that integrate heterogeneous data sources to improve prediction accuracy [goetz2024]. However, limitations remain: knowledge graphs exhibit taxonomic bias toward well-studied organisms (e.g., *E. coli*, *B. subtilis*), and metabolic models may not capture non-genomic factors (quorum sensing, biofilm formation, pH effects). Future iterations should incorporate phenotypic databases (TraDIS, KEIO collections) and environmental context (temperature, osmolarity, oxygen tension) to improve recommendation specificity.

### Experimental Design Efficiency Gains

The MaxPro optimal blocking design achieves a 99.6% reduction in experimental burden relative to a full factorial of six factors at five levels (69 unique conditions vs. 15,625), translating into roughly two orders of magnitude in reagent and labor cost. Beyond cost, the projection-uniformity criterion provides balanced coverage of all lower-dimensional subspaces, which random sampling cannot guarantee and which is critical for response surface and Pareto analyses where ingredient-pair interactions carry the biological signal [joseph2015maxpro]. The blocking layer further controls plate-to-plate confounders (inoculum density, liquid-handler variance) by distributing high- and low-concentration runs evenly across plates, supporting plate-as-random-effect models that absorb technical variance [kasalicky2025, henson2020culturing]. Compared with one-factor-at-a-time screens of the same factor budget, MaxPro retains the power to detect synergistic combinations (e.g., the methanol × PQQ interaction discussed below) that OFAT cannot expose [viana2021rsm].

### Comparison with Agentic AI Co-Scientists

Scope of the agentic contribution. Our retrospective Det-MD ablation (see Results, “Retrospective comparison to a deterministic literature/MediaDive baseline”) locates the agentic contribution narrowly. A non-agentic literature/MediaDive pipeline applied to the same MP-base and the same MaxPro generator recovers four of the six MGA-chosen factors (phosphate, (NH₄)₂SO₄, CoCl₂, methanol) purely from organism-frequency ranking. The two factors that survive only under the agentic pipeline are PQQ and succinate, both of whose supporting evidence lives in literature, genome annotation, and curated FACTS rather than in any media catalog. Det-MD’s mistaken positives — NaCl and glucose — illustrate the converse: frequency-only ranking misfires for AM1 (marine-strain bias for NaCl, non-methylotrophic carbon-source default for glucose), which the agentic layer’s organism-aware judgement correctly avoids. The agentic contribution of MicroGrowAgents is therefore not autonomous factor discovery but a narrower triad: surfacing evidence that lives outside MediaDive, blocking frequency artefacts that don’t match the target biology, and recording the audit trail that lets a reviewer re-derive each decision. This narrowing is the explicit scope we ask Patterns readers to evaluate; the comparison to other agentic co-scientists below is framed accordingly.

A growing family of agentic AI co-scientists now runs autonomous hypothesis–test cycles over user-supplied data, including domain-agnostic systems such as OpenScientist [reese2026openscientist] (an MCP-native loop that iterates over biomedical datasets via sandboxed Python execution), tool-augmented chemistry agents such as ChemCrow [bran2024chemcrow] and the robot-coupled Coscientist [boiko2023coscientist], and open-ended discovery loops such as the AI Scientist [lu2024aiscientist]. OpenScientist is the most directly comparable prime example: it targets the same data-to-hypothesis, wet-lab-adjacent problem MicroGrowAgents addresses, but domain-agnostically. As a class these co-scientists exchange broader domain coverage for less per-domain encoding (we do not attempt to quantify this ratio in either direction), and the contrast clarifies where MicroGrowAgents’ design choices pay off and where they leave headroom.

MicroGrowAgents’ principal advantages over a general-purpose co-scientist loop arise from three architectural commitments. The first is domain-knowledge encoding: curated facts on MxaF/XoxF biochemistry, the PQQ operon, the ethylmalonyl-CoA cycle, an EC-stamped *M. extorquens* AM1 genome-scale model, and per-ingredient DOI provenance let MicroGrowAgents reason about plausibility before an experiment is ever proposed, whereas a domain-agnostic loop must rediscover or hallucinate this biology each run. The second is validation rigor: schema-validated I/O, deterministic seeds, chemical-property checks, genome-scale pathway-gap feasibility checks, and Monte-Carlo Pareto-stability analysis form a layered cascade that is structurally stronger than a single LLM hypothesis-test pass with literature cross-check. The third is DBTL closure: MicroGrowAgents’ recommendations land in actual plate layouts that are executed at the bench and returned to the same analysis pipeline, whereas these co-scientists generally have no concept of physical execution beyond the LLM sandbox — Coscientist, which drives a robotic chemistry platform, is the partial exception. The conscious design choice to keep biology as schema and curated facts rather than as prompt-time context is what makes this rigor auditable for an organism-specific platform [orth2010, price2025kgmicrobe]. These systems also deliver a different kind of transparency than MicroGrowAgents. A co-scientist’s execution log — exemplified by OpenScientist’s MCP-native trace — is strong process transparency: any user can re-trace which tool ran when, with what arguments. MicroGrowAgents’ YAML manifests, LinkML schemas, and per-ingredient DOIs instead deliver evidence transparency, letting a reviewer trace the empirical or literature warrant for any single recommendation back to its source. The two are complementary rather than interchangeable: general-purpose co-scientists optimize for breadth of autonomous action, MicroGrowAgents for depth of justifiable evidence.

The most significant drawbacks are exposure gaps rather than capability gaps. MicroGrowAgents currently exposes its knowledge through Python imports automation recipies in the justfile format rather than as a Model Context Protocol server — the interface most of these co-scientists adopt — which limits portability to other harnesses and is the primary item on our integration roadmap; wrapping query_ingredient, recent_round_winners, lookup_doi, and predict_growth as MCP tools is a tractable next step that preserves all existing agent logic. MicroGrowAgents also omits the provider-agnostic LLM abstraction and per-job spend ledger that systems such as OpenScientist ship, but this is a deliberate non-adoption rather than a capability gap: under a flat-rate subscription harness the marginal per-call cost is essentially zero, so a spend ledger would read zero and provider switching would mean leaving the subscription. The capability would become load-bearing only if a workload moved to per-token API billing (cloud batch or fan-out parallel analysis), at which point it extends cleanly from the existing per-session manifests. Finally, MicroGrowAgents has not yet run a randomized-negative-control design arm of the kind some co-scientists report (OpenScientist demonstrated one across four biomedical case studies). The pipeline is, however, already validated experimentally rather than only in silico: its designs were executed at the bench across two DBTL rounds, the leading composite candidate grew substantially more biomass and faster than the standard base medium, and the held-out benchmark above shows that the six designed factors quantitatively predict Round-2 growth. In a cultivation context, experimentally confirming a system-generated design that outperforms the standard medium is itself a significant result; the remaining gaps are specific rather than wholesale — a randomized or reasoning-ablated control arm to show the system rejects noise, and same-platform confirmation of the chemistry-confounded apparent residual-Nd depletion winner.

### Explainability and Auditability vs Fully LLM-Routed Co-Scientists

A practical corollary of the architectural choices above is that MicroGrowAgents is materially easier to explain, audit, and defend against adversarial inputs than a co-scientist whose every analytical step is routed through an LLM. MicroGrowAgents is itself a coding-agent system — it runs on an LLM coding harness and records its agents’ reasoning and tool calls as inspectable JSONL action traces — so, like a co-scientist, its decision process is analyzable after the fact; in this respect it is closer to a co-scientist than to a black box. What distinguishes it is that this agent reasoning sits on top of a deterministic, schema-grounded core rather than driving the load-bearing decisions. The gap therefore rests on four properties that fully LLM-routed co-scientists, including general-purpose sandboxed-Python loops, do not deliver by construction.

#### Knowledge as schema, not as prompt

MicroGrowAgents encodes biology in LinkML schemas, curated FACTS files (e.g., the MxaF/XoxF lanthanide switch, the PQQ operon, the ethylmalonyl-CoA cycle), and an EC-stamped genome-scale model. Each fact is version-controlled, citable to a DOI, and inspectable independently of any LLM call. In LLM-only co-scientists the same biology lives in opaque parametric weights or is injected as transient context, which is faster to set up but cannot be diffed, reviewed, or signed off by a domain expert before the run.

#### Deterministic core, agentic edges

MaxPro generation, optimal blocking, replicate Pareto computation, and Monte-Carlo stability analysis are seeded and reproducible bit-for-bit; agents enter only at the interpretation boundary, not in the path that selects which conditions go on a plate. The recorded input checksum and fixed random seed reproduce the same plate layout on any machine. Pipelines that route every analytical step through a sandboxed LLM forfeit this property because LLM outputs vary across runs even at temperature zero, and the reasoning that drove a particular recommendation is opaque chain-of-thought rather than a typed function of typed inputs.

#### Recordable decision attribution

When MicroGrowAgents nominates MPOB_058 as a reference condition for a subsequent round, the decision traces back to specific MC-Pareto frequencies (0.922 vs 0.677 vs 0.514), specific replicate sigma estimates, and specific ingredient concentrations stored in queryable session manifests and JSONL action logs. Once the source release and Zenodo snapshot are public (see Data and Code Availability), a reviewer should in principle be able to re-derive each recommendation from the manifest without re-running any agent; we treat fresh-clone re-derivation as a verifiable claim only after end-to-end reproduction has been demonstrated from the archived release. In an LLM-only loop the equivalent recommendation collapses to the model’s terminal narrative, verifiable only by costly re-execution under the same model weights.

#### Resilience to indirect prompt injection

Systems that retrieve text from PubMed, web pages, or per-round knowledge-state files and feed it back into the LLM are structurally vulnerable to indirect prompt injection, where instructions embedded in retrieved content silently redirect the agent’s behavior [greshake2023injection]. MicroGrowAgents narrows the attack surface: text entering via literature retrieval, KG-Microbe queries, or experimental logs is consumed as data inside schema-validated fields, not as control flow for the design or analysis pipelines. An injected instruction in a retrieved abstract becomes a string in a LiteratureAgent evidence record; it cannot rewrite a plate layout or relabel a Pareto winner because those stages are deterministic Python with typed schemas, not LLM-routed tool use. The bound is not absolute — the agentic interpretation layer remains exposed and warrants its own input sanitisation — but the blast radius is restricted to summary text rather than the experimental record.

These properties matter most in two settings: regulated environments (pharmaceutical, clinical) where every design decision must be auditable, and long-running DBTL campaigns where a wet-lab outcome must be unambiguously attributable to a specific predicted condition. Explainability and adversarial robustness are the return on MicroGrowAgents’ deliberate narrowness relative to a general co-scientist. MicroGrowAgents is, by design, semi-deterministic: its load-bearing design and analysis steps are reproducible algorithms, while LLM agents contribute evidence aggregation and interpretation at the edges, and every recommendation is checked both by a human in the loop on design selection and experimentally at the bench. This is a deliberate point on the autonomy spectrum rather than an endpoint. Future iterations will widen the agentic envelope — more autonomous exploration of the design space, agent-proposed design sub-arms tested against deterministic baselines, and closed-loop self-improvement in which the system learns from each DBTL round’s outcomes — while retaining the deterministic, auditable core and the human and experimental validation that make its recommendations trustworthy today.

### Limitations and Future Directions

Despite these advantages, MicroGrowAgents exhibits several limitations. First, knowledge graph coverage remains biased toward culturable organisms with sequenced genomes. Unculturable taxa—the primary targets of media optimization—often lack genomic or metabolic pathway annotations, limiting the system’s ability to generate informed recommendations. For example, candidate phyla radiation (CPR) bacteria possess highly reduced genomes with atypical metabolic networks poorly represented in ModelSEED [lewis2021]. Addressing this requires expanding knowledge graphs to include metagenomic data, single-cell genomics, and trait predictions from phylogenetic placement.

Second, metabolic models may not capture all growth requirements. Essential factors such as organic growth factors (vitamins, amino acids, fatty acids), trace elements at nanomolar concentrations, signaling molecules (autoinducers, siderophores), and environmental parameters (pH, redox potential, osmolarity) are often absent from genome-scale reconstructions [orth2010]. Constraint-based feasibility analysis assumes optimal enzyme kinetics and neglects regulatory networks (transcriptional regulation, post-translational modification) that modulate metabolic flux in response to environmental conditions. Integration of kinetic models, transcriptomic data, and phenotypic screens would improve prediction accuracy but increases model complexity and data requirements.

Third, the system requires experimental validation to confirm predictions. While MaxPro designs efficiently explore parameter spaces, they do not eliminate the need for growth experiments. Laboratory constraints—limited throughput, reagent availability, equipment downtime—restrict the number of conditions that can be tested concurrently. For organisms with multi-week doubling times or fastidious cultivation requirements, this bottleneck persists. Future work should integrate active learning strategies that iteratively refine designs based on experimental results [greenman2025activelearning, sun2024advancingal]. After observing growth outcomes for an initial MaxPro design, a Bayesian optimization layer could propose a second-round design targeting regions of parameter space showing promising results, accelerating convergence to optimal formulations.

Fourth, computational cost scales poorly for high-dimensional parameter spaces. The MaxPro optimization algorithm evaluates millions of candidate designs (n_candidates=100,000), each requiring distance calculations across all design points. For problems with > 20 variable ingredients, runtime exceeds practical limits (hours to days) without advanced optimization techniques. Parallelization across GPUs, gradient-based optimization via automatic differentiation, or surrogate models (Gaussian processes, neural networks) could mitigate this scaling challenge but require substantial implementation effort.

Most consequential for the scope of our claims, the wet-lab evidence here comes from a single organism (M. extorquens AM1) and a single design executed over two cultivation rounds; the biological results therefore demonstrate depth — calibrated, uncertainty-aware ranking with paired controls — rather than cross-organism breadth. We accordingly frame the generalizable contribution as the reusable, evidence-grounded design-and-provenance infrastructure rather than any organism-specific finding (see The Bigger Picture), and treat transfer to additional organisms and chemistries as the necessary next validation step. Within Round-2, leave-one-condition-out benchmarking already shows that the designed factors carry learnable signal about growth (Supplementary Table S2); establishing that the integrated pipeline outperforms simpler, non-integrated baselines across organisms, however, will require running the same workflow on several taxa with independent ground truth, which we have not yet done.

Future directions include integration with automated liquid handling platforms (Hamilton, Tecan, Opentrons) to enable closed-loop experimentation where agent-designed media formulations are robotically prepared, inoculated, monitored, and analyzed without human intervention [bultelle2024autobiotech, papanek2024automation]. Machine learning models trained on accumulated growth data (thousands to millions of organism-media-growth triplets) could replace or complement FBA for growth prediction, particularly for poorly characterized organisms lacking high-quality genome-scale models. Expansion to additional application domains—extremophile cultivation, synthetic community assembly, fed-batch bioreactor optimization—would demonstrate generalizability and drive community adoption.

### Interpreting the Round-2 Pareto Results

The Round-2 cultivation tests the full agents-to-cultivation path. Computing the deterministic Pareto front from replicate means alone yields three winners, but Monte-Carlo resampling against replicate σ separates them sharply: MPOB_058 sits at frequency 0.922 while MPOB_008 and MPOB_019 fall to 0.677 and 0.514. This stratification guards against a common failure mode of multi-objective screens, where “winners” owe their front-membership to favorable noise on a single axis. A paired-control biology analysis at the t2 endpoint refines this further: MPOB_058 is the composite stability winner but is chemistry-confounded (abiotic drift ≈4× any other Pareto winner), whereas MPOB_008 borderline-passes all three independent filters (MC stability, t2 biology signal, near-clean abiotic baseline) and is the cleaner biological-signal anchor (lower abiotic-drift contribution to its apparent Nd depletion). These prioritized conditions are carried forward into subsequent design–build–test–learn rounds for confirmation and replication, with the cycle iterating as further measurements refine the candidate set. Biologically, the trade-off between MPOB_058 (high biomass, moderate Nd uptake) and MPOB_019 (low biomass, deepest Nd uptake) suggests that biomass and apparent residual-Nd depletion objectives are partly antagonistic under the surveyed regime — a tension MicroGrowAgents will need to navigate explicitly when proposing the next round’s designs. A note on cross-round reproducibility bounds how strongly any single round should be read: because Rounds 1 and 2 executed the same v10 design on the same organism, the 69 matched conditions invited a direct reproducibility test, and condition-level Spearman rank correlation between rounds was effectively zero on both axes (ρ = −0.109 for biomass, ρ = +0.058 for apparent residual-Nd depletion). The dominant cause is a deliberate measurement-platform switch — Round-1 used single-channel 600 nm absorbance and uncalibrated arsenazo, whereas Round-2 used a 144-timepoint Biolog Odin kinetic and Miller-calibrated Nd — so the near-zero correlation reflects measurement-modality change rather than biological drift. We therefore treat Round-2 as the calibration-grade dataset and seed subsequent rounds from Round-2 winners alone, and frame the next round as a confirmation round for the prioritized anchor set.

### Provenance, Portability, and Where MicroGrowAgents Sits Among Existing Frameworks

The reproducibility framing above also clarifies how MicroGrowAgents relates to existing infrastructure for portable scientific computation. Workflow languages (Snakemake [koster2012snakemake], the Common Workflow Language [crusoe2022cwl]) and community-curated pipeline registries (nf-core [ewels2020nfcore]) provide strong process transparency: any user can re-execute the same DAG of computational steps in the same order. RO-Crate [soiland2022rocrate] and FAIR Digital Objects [desmedt2020fdo] add machine-actionable provenance metadata at the artefact level. MicroGrowAgents is complementary rather than redundant to these layers: its YAML session manifests, JSONL action logs, LinkML-validated schemas, and per-ingredient DOI provenance are aimed at evidence-level transparency, letting a reviewer trace why a specific ingredient was added at a specific concentration, not just which scripts ran. A productive integration path is to export each completed MicroGrowAgents campaign as an RO-Crate package whose payload includes the session manifests, the design CSV, the wet-lab data, and the Zenodo-archived code release tag; the workflow-language layer can then drive batch re-execution. Adopting that integration is on the roadmap and would close the deployable-to-other-labs gap noted in the Limitations section.

### Broader Impact

Open-source availability of MicroGrowAgents lowers the cost barrier to systematic media optimization. Commercial media optimization services cost $10,000-$50,000 per organism, creating barriers for academic researchers and small biotechnology companies [sipkema2024]. By releasing MicroGrowAgents under a permissive BSD-3-Clause license with complete documentation, tutorials, and provenance-tracked workflows, we enable any laboratory with Python and standard laboratory equipment to perform systematic media optimization.

Applications span diverse microbiology domains. In biofuel production, optimizing growth media for cellulolytic bacteria and methanogens could reduce fermentation costs by increasing cell yields and product titers. In bioremediation, cultivating organisms capable of degrading recalcitrant pollutants (PCBs, per- and polyfluoroalkyl substances) requires tailored media that support growth while inducing catabolic pathways. In human microbiome research, culturing previously unculturable gut bacteria enables functional studies linking microbial metabolism to host health, potentially revealing therapeutic targets [lewis2021]. In synthetic biology, optimizing production strains for high-value compounds (pharmaceuticals, specialty chemicals, materials) requires media formulations that balance growth and product synthesis.

The system’s YAML-based provenance manifests with per-session input checksums and structured agent action logs are designed to make the reported workflows re-runnable from their recorded inputs, pending end-to-end verification from a fresh clone at submission; this addresses a recurring barrier to independent verification in computational biology. This audit trail proves particularly valuable in regulatory environments (pharmaceutical manufacturing, clinical diagnostics) where data integrity requirements mandate documented evidence of all analysis steps.

## STAR Methods

## Resource availability

### Lead contact

Further information and requests for resources should be directed to and will be fulfilled by the lead contact, Marcin P. Joachimiak (; ORCID [FILL]), Biosystems Data Science Department, Environmental Genomics and Systems Biology Division, Lawrence Berkeley National Laboratory.

### Materials availability

This study did not generate new unique reagents. The *Methylorubrum extorquens AM1:ΔmxaF strain* used in this work is available from the lead contact on reasonable request, subject to a standard material transfer agreement.

### Data and code availability

Raw cultivation data and provenance manifests are deposited at Zenodo and on GitHub; the Key Resources Table below lists deposited-data accessions and DOIs once assigned. Code and data will be made publicly available under the BSD-3-Clause license on publication.

## Key Resources Table

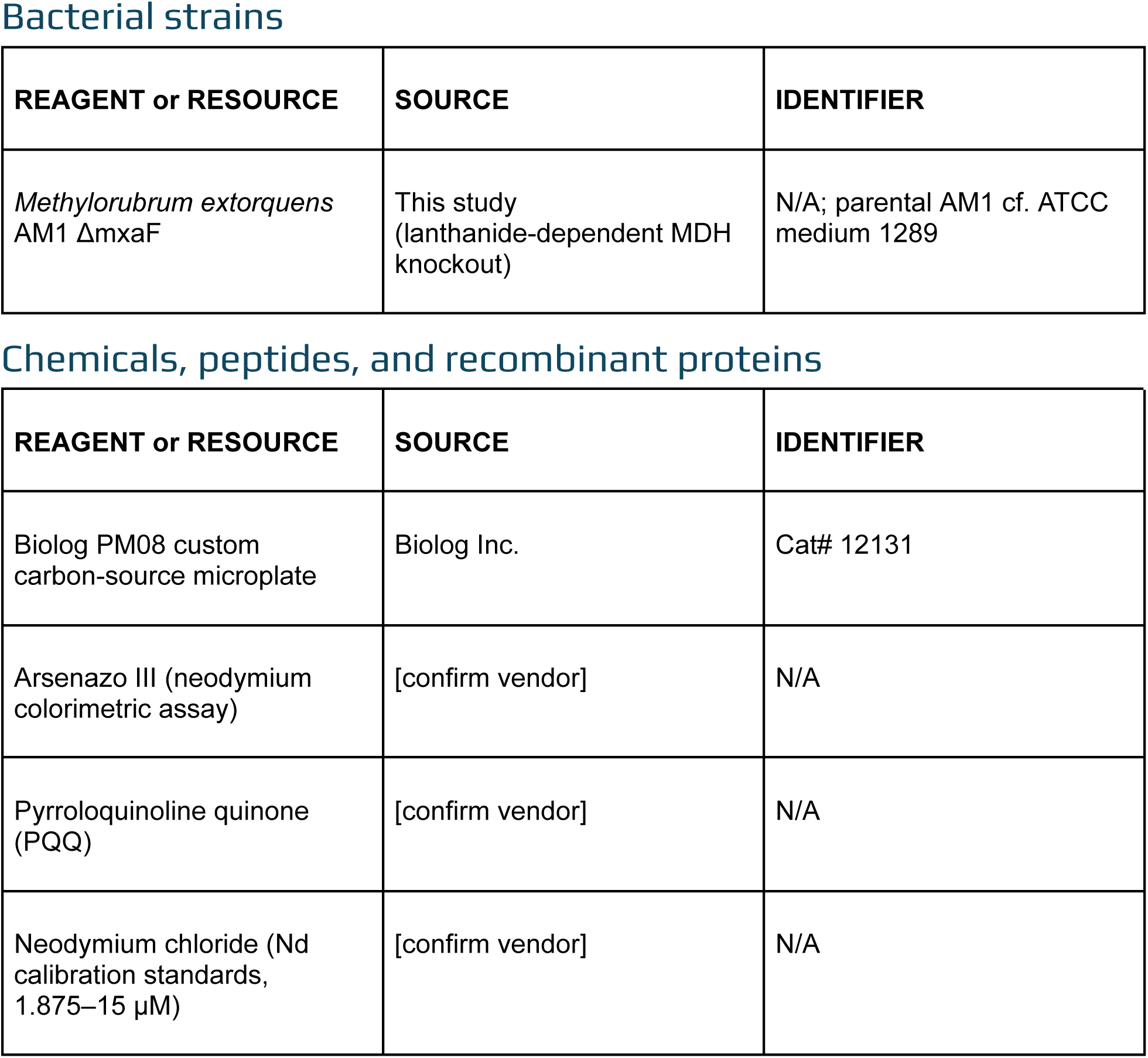
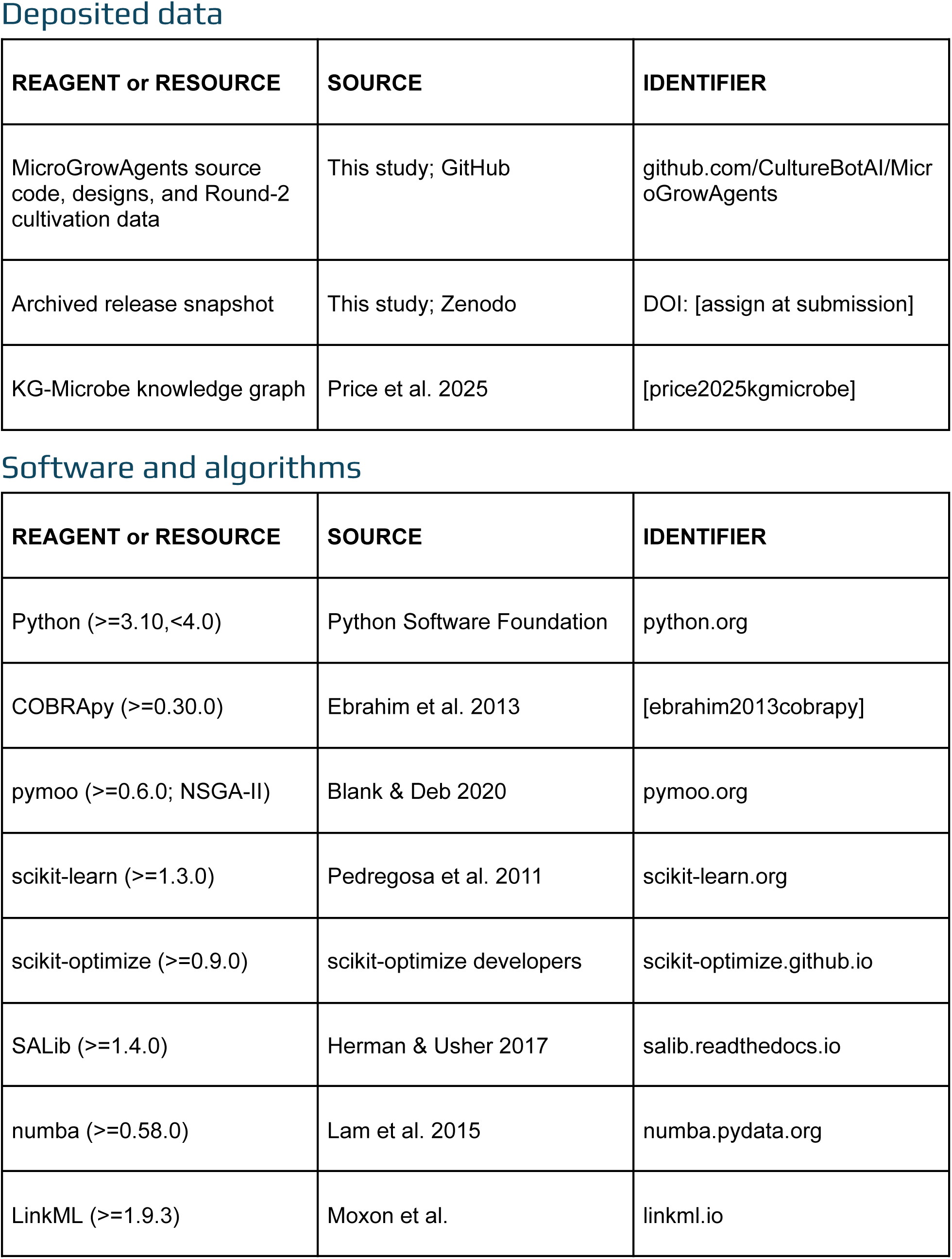
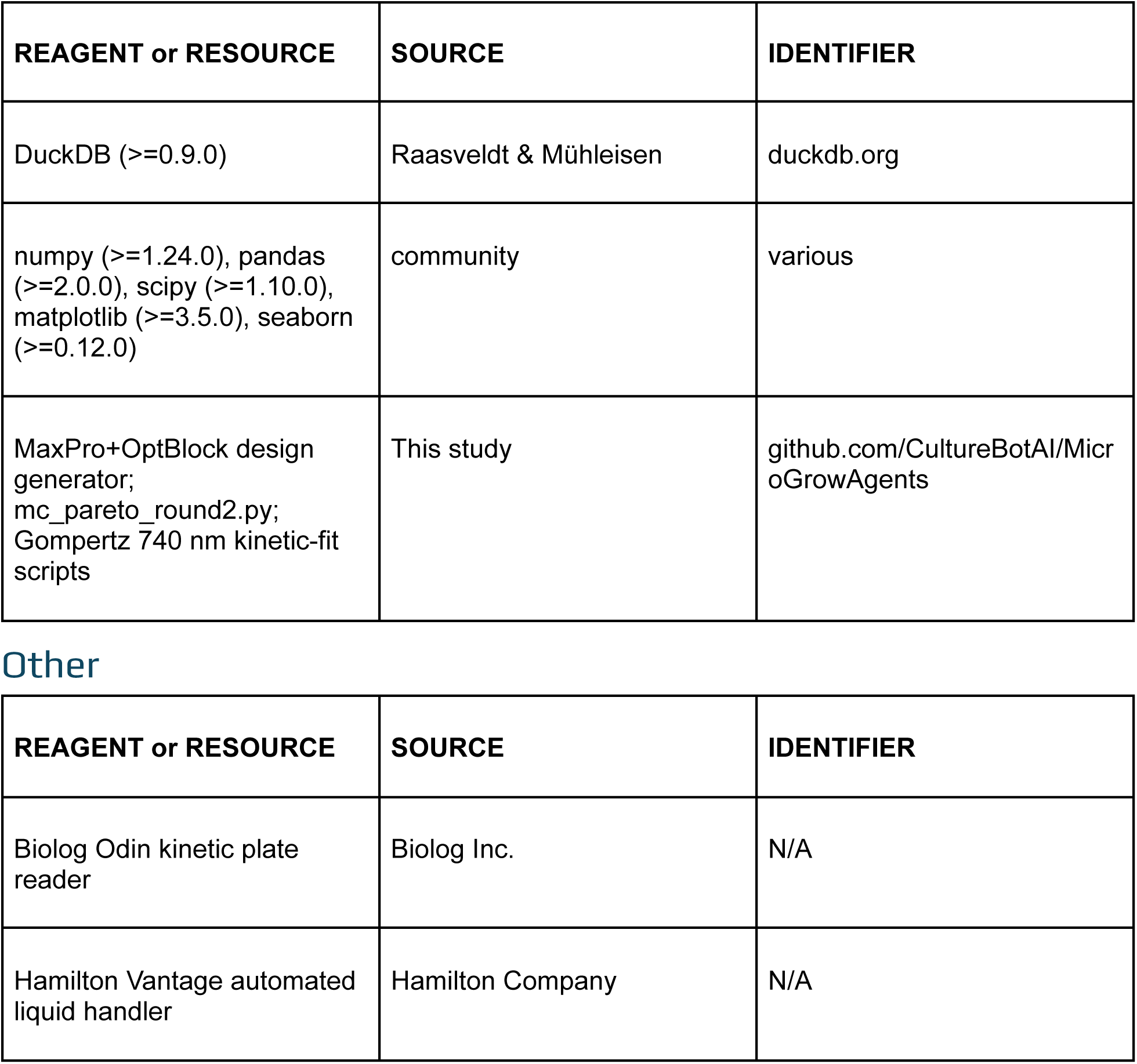

## Experimental model and study participant details

### Culture and Growth of strain

*Methylorubrum extorquens AM1:*△*mxaF* was maintained on MP agar plates (MP medium supplemented with 1.5% agar) and incubated at 30°C; visible colonies appeared within 48 hours. For liquid pre-cultures, a clump of cells from an agar plate was inoculated into 50 mL of MP medium in a 250 mL baffled flask and grown overnight at 30°C with orbital shaking at 200 rpm. The composition of MP medium follows [delaney2019design].

### Growth kinetics in microplate reader

Overnight cultures were harvested by centrifugation and washed three times with carbon-free MP medium (all components except carbon sources) to remove residual nutrients. Washed cells were resuspended to a starting OD600 of 0.1 in each designed medium variation. Assays were conducted in 48-well plates sealed with a gas-permeable membrane. Plates were incubated at 30°C with continuous orbital shaking at 200 rpm in a BioTek microplate reader for 48 hours. Absorbance at 600 nm was recorded at regular intervals to generate full growth curves. Area under the curve (AUC), maximum absorbance, and growth rate were extracted.

### Growth and metabolite measurements in Biolog Odin

Designed media variations were prepared in 96-well deep-well plates (2 mL capacity), and 100 µL of each media design was transferred into a 96-well Biolog PM08 custom carbon-source plate (Biolog Inc., catalog no. 12131) and inoculated with *M. extorquens* AM1 ΔmxaF to a final starting OD600 of 0.1. Plates were then incubated in a Biolog Odin instrument at 30 °C for 48 h. Absorbance at 740 nm (biomass turbidity) was recorded at 20 minute intervals. Each designed condition was represented by four replicate wells; four additional wells containing control media served as plate-level controls.

### Arsenazo III assay for residual neodymium

Residual neodymium (Nd) in the spent medium was quantified by arsenazo III colorimetric assay. At the experimental endpoint, 150 µL of culture from each deep-well plate was transferred to a 250 µL flat-bottom 96-well plate (Corning) and OD600 was measured to confirm cell density. To ensure sufficient clarified supernatant volume and to prevent inadvertent cell pellet carryover, cultures were then transferred to a 96-well PCR plate and centrifuged at 2000 × g for 10 minutes. Supernatant (120 µL) was transferred to a new flat-bottom 96-well plate, and 12 µL of 0.5 M HCl was added to acidify the sample to pH 2.6-2.7. Plates were shaken at 200 rpm for 20 minutes to precipitate PIPES buffer, then centrifuged at 2000 x g for 10 minutes to pellet precipitate. For the colorimetric reaction, 100 µL of citrate-phosphate buffer, 100 µL of clarified supernatant, and 10 µL of arsenazo III reagent were combined in a new flat-bottom 96-well plate, shaken for 10 minutes, and absorbance was measured at 660 nm. Residual Nd concentrations were calculated from a per-plate calibration curve prepared in Row B using Nd standards at known concentrations spanning the expected assay range. Plate-to-plate variation in ctrl_media wells was approximately 5% CV across the experimental plates (ctrl_media OD600 0.205 ± 0.010, n = 16 wells across the four-plate Biolog execution).

## Method details

### Software Architecture

MicroGrowAgents is implemented in Python 3.10+ as a modular agent-based system integrating large language models with biological knowledge bases. The architecture comprises 29 documented agents and 58 documented skills organized into seven functional categories: knowledge and database access, modeling, chemistry and analysis, media design, experimental design, evidence and enrichment, and analysis and interpretation (a broader internal inventory including workflow-composed agents enumerates 32 agents and 63 skills). Each agent is implemented as a Python class inheriting from a base Agent interface that handles prompt construction, API communication with Claude 4 family models [anthropic2025claude], error handling, and result validation. The agents run within a coding-agent harness — a runtime (Claude Code) in which an LLM can read and write files, execute Python and shell tools, and invoke the agent skills — that orchestrates each agent’s tool calls and records every read, query, decision, and code execution as a structured JSONL action log. This harness is what makes the agents’ reasoning and actions inspectable after the fact (see Discussion).

The system uses a skill-based design where agents execute specific capabilities defined as structured prompts with input schemas, execution logic, and output validation. Skills are stored in .claude/skills/ as markdown documents following a standardized template specifying name, description, required inputs, tool permissions, and execution context. This approach enables rapid prototyping and modification of agent behaviors without code changes. Configuration management uses pyproject.toml for dependency specification and YAML manifests for runtime parameters, ensuring reproducible executions across environments.

Testing infrastructure employs pytest with unit tests for individual agent methods, integration tests for multi-agent workflows, and validation of complete media recommendation pipelines. Dependencies include scientific Python stack (numpy, pandas, scipy, scikit-learn), metabolic modeling frameworks (cobra, gemsembler), knowledge graph tooling (KGX, Biolink Model, METPO, LinkML, KG-Registry), visualization libraries (matplotlib, seaborn), and optimization packages (scikit-optimize, pymoo, salib). Performance-critical MaxPro pairwise-distance kernels use numba just-in-time compilation; full optimization of the v10 design (n_candidates = 100,000, 13 iterations) ran to convergence in 106 s on a single CPU core, with single MaxPro evaluations completing in well under one second.

### Agent and Skill Implementation

Agents are organized into seven functional categories. The Knowledge & Database category includes KGReasoningAgent for SPARQL queries against KG-Microbe [price2025kgmicrobe], LiteratureAgent for PubMed/Semantic Scholar searches, SQLAgent for ingredient-property DuckDB queries, and SheetQueryAgent for structured data lookups. Modeling provides GEMsemblerAgent (via COBRApy [ebrahim2013cobrapy]) for genome-scale model reconstruction and flux balance analysis, GenomeFunctionAgent for Bakta-annotated genome interpretation, LanthanideGenesAgent for rare-earth-element metabolism detection, TransporterAgent for nutrient uptake system annotation, and MetabolicSourceAgent for C/N/P/S source compatibility analysis. Chemistry & Analysis includes ChemistryAgent for osmotic property calculation, redox potential prediction, and chemical similarity using RDKit Morgan fingerprints. Media Design implements MediaFormulationAgent for multi-source recommendation synthesis, GenMediaConcAgent for ML-based concentration prediction using scikit-learn random-forest regressors, and CofactorMediaAgent for vitamin/cofactor requirement analysis. Experimental Design hosts MaxProOptBlockAgent for projection-uniform optimal blocking designs and IngredientCooccurrenceAgent for TF-IDF-like statistical analysis of ingredient co-occurrence across the MediaDive corpus (3,327 media formulations), which scores pairwise associations as TF (co-occurrence count / media containing the first ingredient) times IDF (log(total media / co-occurrence count)) so that pairs that occur frequently together but are not ubiquitous (e.g., PQQ with neodymium) surface as candidate evidence for ingredient recommendations rather than ubiquitous pairings (e.g., glucose with ammonium). Evidence & Enrichment includes PDFEvidenceExtractor, EvidenceExtractionOrchestrator, IngredientEffectsEnrichmentAgent, and CSVAllDOIsEnrichmentAgent for evidence harvesting and DOI-linked enrichment. Analysis & Interpretation provides AnalogyReasoningAgent for cross-organism inference and ReconcileAgent for multi-source consensus.

Each skill defines allowed tools (Read, Write, Bash, Grep, Glob) and execution context (fork vs. maintain). Fork context creates isolated conversation threads for complex multi-step workflows, while maintain context preserves conversation history for iterative refinement. Prompt engineering follows structured templates: system instructions defining agent role and capabilities, context section providing relevant data (genome annotations, literature evidence, media databases), instruction section specifying the task, constraints section defining validation criteria, and output section specifying format requirements (YAML, JSON, TSV). Prompts incorporate few-shot examples for complex reasoning tasks and chain-of-thought templates for multi-step decision processes.

Action provenance is recorded in YAML manifests stored in .claude/provenance/sessions/ directories. Each manifest captures session metadata (timestamp, model version, goals), action sequence (tool calls, parameters, results), files affected (paths, checksums, sizes), decisions made (rationale, alternatives considered), and validation results (schema compliance, data quality checks). This enables workflow reproducibility and audit-trail generation.

### Data Source Integration

MicroGrowAgents integrates eight external knowledge bases to inform media recommendations. KG-Microbe [price2025kgmicrobe] provides organism taxonomy (864,363 species from GTDB [parks2024], LPSN, NCBI), metabolic pathways, growth conditions, and media-organism associations via SPARQL endpoint queries. SPARQL queries target specific organism taxa (genus Methylorubrum), metabolic capabilities (methylotrophy, C1 metabolism), and documented growth media. ModelSEED Biochemistry Database [seif2021modelseed] supplies reaction stoichiometry, compound structures, and metabolic pathway templates accessed via REST API with caching to minimize redundant requests.

PubChem REST API retrieves chemical properties (molecular weight, LogP, solubility, SMILES) for ingredient validation and alternative suggestions. ChEBI ontology [hastings2016] provides hierarchical chemical classification, enabling functional group-based searches and ingredient substitution logic. PaperBLAST integration mines literature evidence linking genes to metabolic functions, implemented via local installation of GapMind tools and cached result databases. LPSN taxonomy integration [parks2024] validates organism nomenclature and retrieves phylogenetic context for analogical reasoning.

Data integration workflow: (1) organism name normalization via fuzzy string matching against LPSN/NCBI taxonomy, (2) SPARQL query construction with genus/species filters, (3) API requests with exponential backoff retry logic, (4) response parsing with schema validation, (5) result caching in local DuckDB database for offline access. All external data retrievals are logged with timestamps, checksums, and API versions in provenance manifests. Cached data undergoes periodic refresh checks comparing local checksums against remote versions, triggering re-downloads when mismatches occur.

## Quantification and statistical analysis

All analyses below use n=4 replicate wells per designed condition, with four control wells per plate (16 total across the four-plate Biolog execution) supporting per-plate normalization. Primary endpoints are reported at the t2 plate-reader timepoint: abiotic-correction diagnostics indicated that t2 captured biomass signals before precipitation artifacts began to dominate at t3, making it the most representative of biological signal. Significance testing uses two-sample t-tests with Bonferroni correction for multiple comparisons (α = 0.05 / n_conditions). Replicate outliers were identified via hierarchical clustering (Ward linkage, Euclidean distance) and removed when located more than three standard deviations from the cluster centroid; this retained >95% of wells. Replicate-level uncertainty is propagated downstream via the Monte-Carlo Pareto frontier procedure described below. To test how much of the Round-2 growth outcome is predictable from the six designed factors, we ran leave-one-condition-out cross-validation over the 70 conditions: each condition was held out in turn, a model (random forest, gradient boosting, or Gaussian process; a constant global-mean predictor as the floor) was trained on the remaining 69 conditions, and we scored held-out R², mean absolute error, Spearman rank correlation, and Precision@5 (recovery of the true five highest-AUC conditions). A stoichiometric flux-balance baseline on the AM1 consensus genome-scale model (cobrapy) was evaluated by scaling the variable media-exchange bounds with each condition’s concentrations. We did not cross-validate across rounds because Round-1 and Round-2 used incompatible measurement modalities; the analysis script and outputs are released with the repository.

### Experimental Design Generation

MaxPro optimal blocking design [joseph2015maxpro, yang2024] generates space-filling experimental conditions with minimized batch effects. The design process begins with ingredient selection: the baseline MP medium comprises 20 annotated ingredients (excluding buffer) partitioned into fixed (held constant across conditions) and variable (optimized) components. For *M. extorquens* AM1 lanthanide depletion study, six factors were selected as designed variables — Total_Phosphate, (NH₄)₂SO₄, CoCl₂, Succinate, Methanol, and PQQ — on the basis of literature evidence, genome-scale pathway-gap analysis, and cofactor requirements; remaining ingredients were fixed at MP-standard concentrations, and CaCl₂, citrate, biotin, and thiamine were excluded by design choice for this campaign. Latin Hypercube Sampling (LHS) [mckay2000] generates candidate designs by partitioning each ingredient’s concentration range into *n* equal-probability intervals and randomly sampling one value per interval, ensuring uniform coverage across the parameter space.

The MaxPro criterion maximizes the minimum projection distance:

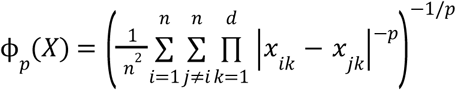

where *X* is the *n* × *d* design matrix (*n* runs, *d* ingredients), *x_ik_* is the normalized concentration of ingredient *k* in run *i*, and *p* = 15 (default parameter). Higher ϕ_p_ values indicate better space-filling. Optimization employs coordinate exchange algorithm: iterative replacement of individual design points with candidates maximizing ϕ_p_, repeated until convergence (change < 10^-6^) or maximum iterations (10,000) reached.

Blocking assigns conditions to experimental plates minimizing correlation between plate identity and ingredient concentrations. Given 69 unique conditions replicated 4 times (276 runs) distributed across 3 plates (92 runs per plate), blocking variables are selected as the 3 highest-variance ingredients. Block assignment uses optimal partition algorithm minimizing within-block variance:

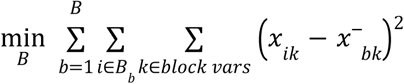

Where *B* = {*B*_1_, …, *B_B_*} partitions runs into *B* = 3 blocks, *x*^−^*_bk_* is the mean of blocking variable *k* in block *b*. This ensures each plate observes similar ranges of critical ingredients, reducing plate-to-plate confounding.

Control samples include 4 baseline MP medium wells per plate (12 total), allocated to randomized well positions. Additional controls (sterile medium, uninoculated medium) are handled via split-plate protocol: prepare 1 mL of each condition, split into 2×0.5 mL aliquots, inoculate only one set. This approach doubles effective sample size while maintaining independent negative controls.

Concentration range selection employs three strategies: (1) literature-derived ranges from MediaDive database queries (median ± 2 standard deviations), (2) stoichiometric estimates of nutrient demand from cell-composition and pathway requirements, (3) expert constraints (e.g., methanol maximum 150 mM to avoid toxicity, minimum 10 mM to support growth). Final ranges are validated via sensitivity analysis sweeping concentrations in 10% increments and predicting growth impacts using trained ML models.

The genome annotations and curated organism FACTS sheets determined which cofactors and nutrients were eligible to enter the design as variable factors rather than fixed background ingredients; their concentration ranges were set by the literature/MediaDive, stoichiometric, and expert strategies above and refined across experimental rounds by sensitivity analysis of prior-round data, not from genome-derived essentiality scores.

Design outputs include: design_matrix_v10.csv (276 runs × 15 ingredient concentrations), block_assignments_v10.csv (run-to-plate mapping), selected_conditions_v10_maxprooptblock_long.csv (unique conditions with metadata), stock solution preparation protocols, Hamilton liquid handler transfer instructions, and data collection templates. All outputs undergo LinkML schema validation ensuring compliance with internal data standards.

### Response Surface Modeling and Analysis

Growth improvement analysis compares experimental optical density (OD600) measurements against baseline controls. Growth improvement is calculated as:

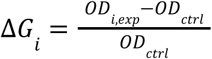

where *OD_i,exp_* is the final OD600 for experimental condition *i* (mean of 4 replicates) and *OD_ctrl_* is the control plate mean. Statistical significance assessed via two-sample t-tests with Bonferroni correction for multiple comparisons (α = 0. 05/*n_conditions_*).

Response surface modeling employs radial basis function (RBF) interpolation [regis2007rbf] to predict growth as a continuous function of ingredient concentrations. RBF model:

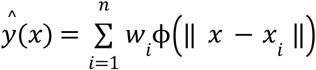

where *x* is the ingredient concentration vector, *x_i_* are the experimental design points, ϕ(*r*) = *exp*(*-r*^2^/(2σ^2^)) is the Gaussian kernel (σ = 0. 5), and weights *w_i_* are solved via least squares fitting to observed growth data. Model validation uses 5-fold cross-validation, reporting mean absolute error (MAE) and *R*^2^.

Interaction effects identified via ANOVA decomposition separating main effects (single ingredient contributions) and two-way interactions (synergistic/antagonistic ingredient pairs). Significant interactions (*p* < 0. 01) are visualized as contour plots showing predicted growth across concentration gradients. Clustering analysis groups experimental replicates via hierarchical clustering (Ward linkage, Euclidean distance) to identify outliers (replicates > 3 standard deviations from cluster centroid). Outlier removal improves model fit while retaining > 95% of data points.

Optimization identifies concentration combinations maximizing predicted growth via multi-objective evolutionary algorithm (NSGA-II) with objectives: maximize growth, minimize ingredient cost, minimize preparation complexity. Pareto-optimal solutions are ranked by desirability scores combining weighted objectives, producing candidate formulations for iterative experimental validation.

### Monte-Carlo Pareto Frontier Analysis

To identify multi-objective optima while explicitly accounting for replicate-level uncertainty, we perform a Monte-Carlo Pareto frontier analysis (scripts/mc_pareto_round2.py) over two objectives: maximize biomass turbidity at 740 nm and minimize residual Nd concentration in spent medium (apparent residual-Nd depletion by arsenazo III colorimetric assay). Endpoint measurements are taken at the t2 plate-reader timepoint, selected after abiotic-correction diagnostics indicated t2 captured biological signal before precipitation artifacts dominate at t3.

For each condition *c* we compute the replicate mean *x*^−^*_c,k_* and standard deviation σ ^*_c,k_* for each of the two response variables *k* ∈ {*OD*600, *Abs*590, *Nd*}. We then draw *N* = 1000 Monte-Carlo samples; each sample independently draws a perturbed coordinate 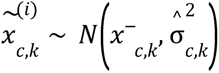 condition per axis, recomputes the two-objective Pareto front p^(i)^, and records front membership. The Pareto-membership frequency for condition *c* is

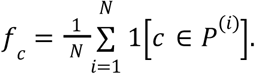

Conditions are classified into four stability tiers: stable (*f*_c_ ≥ 0. 8), borderline (0. 5 ≤ *f*_c_ < 0. 8), rare (0. 1 ≤ *f*_c_ < 0. 5), and off-frontier (*f*_c_ < 0. 1). The stable tier identifies anchor conditions for subsequent DBTL rounds; the borderline tier flags conditions whose front membership warrants additional replication before commitment. The deterministic Pareto front, computed from replicate means alone, is reported alongside MC frequencies for cross-comparison. Per-axis errors are treated as independent across the two objectives.

### Metabolic Modeling Integration

Genome-scale metabolic models (GEMs) for *M. extorquens* AM1 were reconstructed using COBRApy [ebrahim2013cobrapy] and ModelSEED templates [seif2021modelseed]. The reconstruction workflow includes: (1) gene annotation using Bakta, (2) mapping gene products to biochemical reactions via EC numbers and sequence similarity, (3) biomass reaction formulation based on cell composition measurements, (4) pathway-completeness (gap) and auxotrophy analysis to flag missing biosynthetic capabilities. FBA solves:

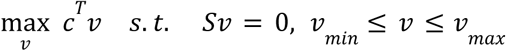

where *v* is the flux vector, *c* is the biomass objective coefficient, *S* is the stoichiometric matrix, and bounds *v_min_*, *v_max_* constrain uptake/secretion rates.

Gap-filling identifies missing reactions preventing biomass production under specified media conditions. Gap-filling optimization:

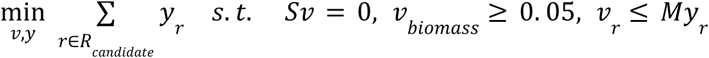

where *R_candidate_* are candidate reactions from universal biochemistry database, *y*_r_ ∈ {0, 1} indicates reaction inclusion, *M* is a large constant, and minimum biomass flux ensures growth feasibility. Gap-filled reactions are reviewed for genomic evidence (homologous genes, domain predictions) before inclusion.

As a general capability — not applied quantitatively to strain AM1, whose available genome-scale reconstructions do not support methylotrophic growth (see Results) — simulated media formulations would constrain uptake fluxes to proposed ingredient concentrations to estimate growth:

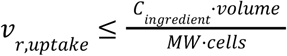

where *C_ingredient_* is the concentration (mM), volume is the culture volume (mL), MW is molecular weight (g/mol), and cells is the estimated cell count at stationary phase. Genome-scale pathway-gap and auxotrophy analysis flagged candidate biosynthetic gaps for review (for example, an initial partial-biosynthesis call for cobalamin that curated facts later corrected; see Results) and hallmark methylotrophy pathways, informing which ingredients were treated as fixed must-haves versus variable factors. We did not use quantitative FBA growth prediction to filter the candidate pool or set the factor ranges, because the available *M. extorquens* AM1 genome-scale reconstructions do not support methylotrophic growth (see Results); the variable-factor ranges were literature- and MediaDive-derived.

## Data and Code Availability

Source code will be released publicly at https://github.com/CultureBotAI/MicroGrowAgents under the BSD-3-Clause license concurrent with the publication of this manuscript, with an archival snapshot deposited at the same time via a Zenodo DOI tied to a tagged release. Installation requires Python 3.10+ and proceeds from a clone of the tagged release (pip install git+https://github.com/CultureBotAI/MicroGrowAgents.git@<release-tag>; the exact tag will be fixed at submission). Complete documentation is hosted at https://CultureBotAI.github.io/MicroGrowAgents. All experimental designs, ingredient databases, cofactor mappings, and analysis outputs are included in the repository under data/ directories. The Round-2 cultivation data, Monte-Carlo Pareto analysis scripts (scripts/mc_pareto_round2.py, scripts/three_way_pareto_round2.py), and resulting tables and figures used in this manuscript are released under outputs/round2_mc_pareto/ and outputs/round2_3way_pareto/. Provenance tracking records for reproducibility are maintained in .claude/provenance/sessions/. System requirements and dependencies are specified in pyproject.toml.

